# Comparative neurotranscriptomics reveal widespread species differences associated with bonding

**DOI:** 10.1101/2020.12.07.415463

**Authors:** Joel A Tripp, Alejandro Berrio, Lisa A McGraw, Mikhail Matz, Jamie K Davis, Kiyoshi Inoue, James W Thomas, Larry J Young, Steven M Phelps

## Abstract

**Background:** Pair bonding with a reproductive partner is rare among mammals but is an important feature of human social behavior. Decades of research on monogamous prairie voles (*Microtus ochrogaster*), along with comparative studies using the related non-bonding meadow vole (*M. pennsylvanicus*), have revealed many of the neural and molecular mechanisms necessary for pair-bond formation in that species. However, these studies have largely focused on just a few neuromodulatory systems. To test the hypothesis that neural gene expression differences underlie differential capacities to bond, we performed RNA-sequencing on tissue from three brain regions important for bonding and other social behaviors across bond-forming prairie voles and non-bonding meadow voles. We examined gene expression in the amygdala, hypothalamus, and combined ventral pallidum/nucleus accumbens in virgins and at three time points after mating to understand species differences in gene expression at baseline, in response to mating, and during bond formation.

**Results:** We first identified species and brain region as the factors most strongly associated with gene expression in our samples. Next, we found gene categories related to cell structure, translation and metabolism that differed in expression across species in virgins, as well as categories associated with cell structure, synaptic and neuroendocrine signaling, and transcription and translation that varied among the focal regions in our study. Additionally, we identified genes that were differentially expressed across species after mating in each of our regions of interest. These include genes involved in regulating transcription, neuron structure, and synaptic plasticity. Finally, we identified modules of co-regulated genes that were strongly correlated with brain region in both species, and modules that were correlated with post-mating time points in prairie voles but not meadow voles.

**Conclusions:** These results reinforce the importance of pre-mating differences that confer the ability to form pair bonds in prairie voles but not promiscuous species such as meadow voles. Gene ontology analysis supports the hypothesis that pair-bond formation involves transcriptional regulation, and changes in neuronal structure. Together, our results expand knowledge of the genes involved in the pair bonding process and open new avenues of research in the molecular mechanisms of bond formation.

## Background

The profound social attachments humans form with one another are a defining feature of our species. While many species develop parent-offspring or mate pair bonds, these attachments are especially strong and long-lasting in humans, particularly those between reproductive partners (1). In fact, the social bonds formed by human partners are quite uncommon among mammals: only 9% of mammalian species display social monogamy, including just 6% of rodents, which are often used as model species to investigate the neural basis of behavior (2). Importantly, commonly used laboratory species including mice and rats are not monogamous and lack the ability to form lasting pair bonds. A better understanding of the mechanisms underlying the formation of partner bonds requires use of a species that forms such bonds.

The voles of the genus *Microtus* (Fig. 1a, b) have been developed into powerful models for better understanding the neural mechanisms underlying attachment and pair bonding (3). Typically, the focal species of these studies is the prairie vole (*Microtus ochrogaster*, Fig. 1a), which has become famous for the tendency for males and females to form long-lasting, socially monogamous bonds. Field studies of prairie vole space use and behavior demonstrated a socially monogamous mating system (4, 5) (though extra-pair fertilization does occur and some individuals adopt more promiscuous mating tactics (6, 7)). In the lab, receptive virgin prairie voles will mate with novel opposite-sex conspecifics and, over a period of several hours of mating and other affiliative behaviors, will form a pair bond characterized by selective affiliation and increased aggression towards intruders (4,5,8,9).

**Fig. 1.**
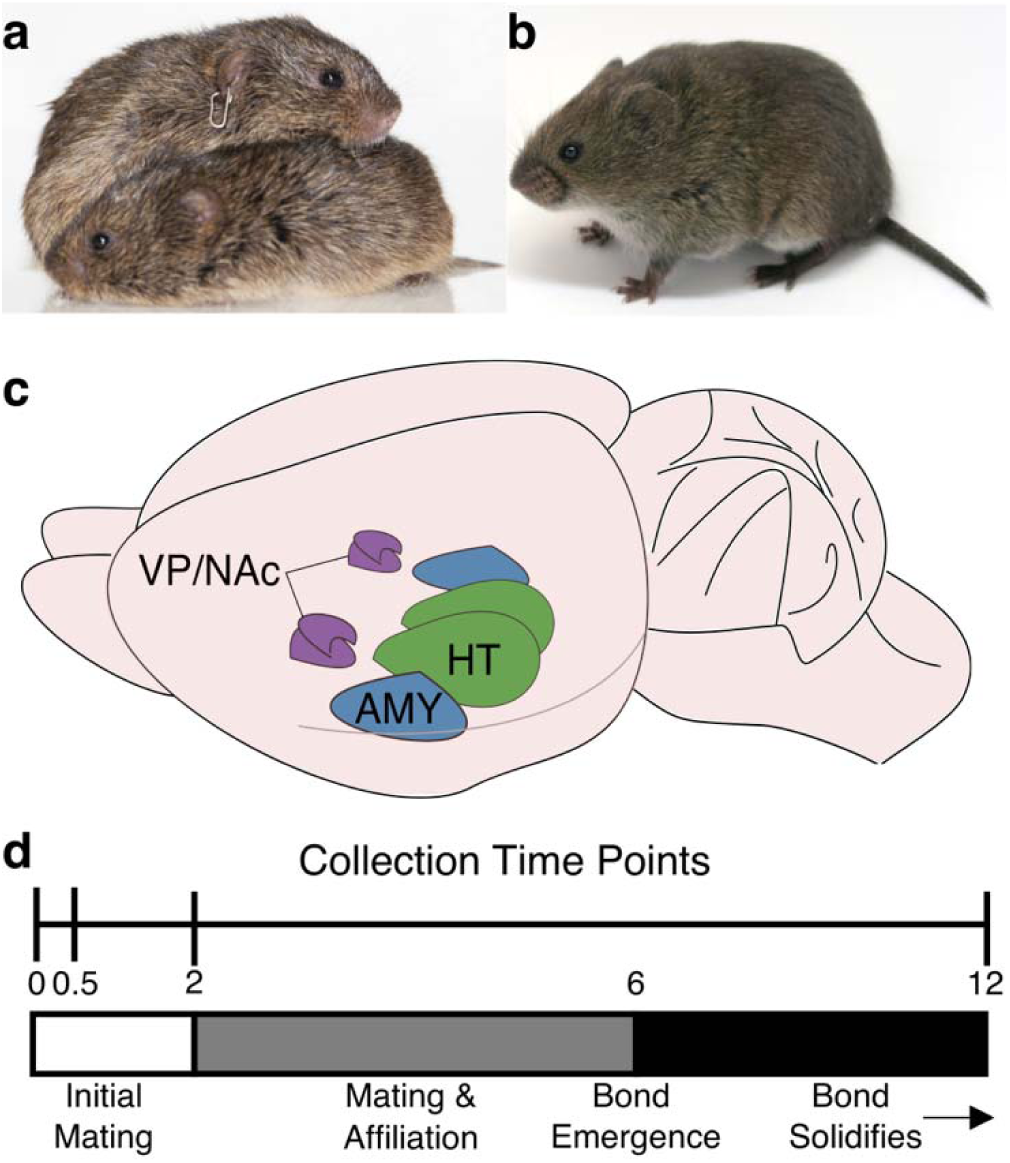
Experiment overview. **a.** Pair of prairie voles (*Microtus ochrogaster*) huddling (Photo credit: Aubrey M. Kelly) **b.** A single adult meadow vole (*Microtus pennsylvanicus*) (Photo credit: Beery Lab) **c.** Drawing of vole brain. Regions of interest for this study are highlighted, including the amygdala (AMY, blue), hypothalamus (HT, green), and ventral pallidum and nucleus accumbens (VP/NAc, purple). **d.** Above: Experiment Timeline. Virgin animals (time 0) were collected prior to mating. Mated animals were collected at either 0.5, 2, or 12 hr after first intromission. Below: Timeline of prairie vole bond formation. Unfamiliar opposite sex conspecifics will mate shortly after introduction. Bonds become detectable at 6 hr after onset of mating and strengthen up to 24 hr after mating onset (8, 9).

Prairie voles are often contrasted with closely related species which do not form these bonds (8,10–14), including the meadow vole (*Microtus pennsylvanicus*, Fig. 1b). Field studies of meadow voles are consistent with a promiscuous mating system and in the lab, they do not show the selective affiliation with mating partners characteristic of pair bonds (12,15,16). These comparative studies have been particularly useful in uncovering the features of the prairie vole brain that facilitate bond formation. These include neuroendocrine signaling in the brain (particularly via the nonapeptides oxytocin [OT] and arginine-vasopressin [AVP]) which conveys social salience, mechanisms of social recognition and memory, and dopaminergic reward pathways (17, 18). Current hypotheses propose that pair bonding results from synaptic plasticity that links the neural encoding of partner cues with the reward system, leading to persistent reinforcement of the partner that results in selective affiliation (17).

While significant progress has been made in understanding the neural mechanisms underlying vole pair-bond formation, previous studies have mainly focused on a relatively small number of neuromodulatory pathways, including the nonapeptides and dopamine reward systems mentioned above, as well as opioid signaling in the brain (17, 18). In order to identify new genes that may be playing a significant role in bond formation, it is necessary to take a broader perspective and examine changes to global gene expression in regions with a known role in bonding.

In this study, we used a comparative approach, taking advantage of the close evolutionary relationship (19) and stark contrast in bonding behavior between prairie and meadow voles to better understand the molecular mechanisms that support pair-bond formation and identify novel candidate genes for future study. Our study focused on three regions (Fig. 1c): the amygdala (AMY), hypothalamus (HT), and a region inclusive of the ventral pallidum and nucleus accumbens (VP/NAc). Each of these regions is proposed to have a critical function in the development of a pair bond (17). The rodent AMY receives olfactory information critical for social recognition (20, 21), encodes valence (22), and plays an important role in social memory (23, 24). The HT contains several nuclei that are involved in regulating social behavior, including populations that produce the neuropeptides OT and AVP, which signal social salience, influence social memory, and regulate related behaviors (12,25–28). Finally, signals of social salience converge with dopaminergic signaling from the ventral tegmental area in the NAc, resulting in disinhibition of the VP, which influences behavioral responses to rewarding stimuli (29–32). While several other regions have important functions in the development and maintenance of pair bonds (17, 18), our focal regions represent key nodes involved in critical aspects of bonding: social context, social memory, and reward.

We used RNA-sequencing to quantify gene expression before and at three time points (0.5, 2, and 12 hr) after mating (Fig. 1d) in each of these regions across both prairie and meadow voles. By observing changes in gene expression in these regions across time relative to mating and across pair-bonding and non-bonding species, we sought to identify genes involved in the process of pair-bond formation, either by conferring the capacity to bond before mating or changing their expression to support bonding after mating. We find gene categories that differ across species at baseline (pre-mating) and across brain regions, as well as genes that differ in expression post-mating between prairie and meadow voles. Further, using gene network analysis, we identify gene modules that have expression patterns strongly related to specific brain regions, as well as modules that change in expression in response to mating. Together, these results provide further support for the current models of mammalian pair bonding and provide additional candidate genes that may underlie the formation of monogamous bonds.

## Results

In order to identify the factors (e.g. species, brain region, time relative to mating) most strongly associated with gene expression in our samples, we first performed hierarchical clustering based on Poisson distance (33), followed by principal component analysis (PCA). In both cases, the primary factor influencing gene expression was species, followed by brain region (Fig. 2). Hierarchical clustering based on Poisson distance resulted in a tree with two major branches representing samples from prairie and meadow voles. These branches were then split corresponding to brain region. In prairie voles, AMY and VP/NAc clustered more closely to each other, while in meadow voles AMY and HT were more closely clustered (Fig. 2a). This result was also supported by PCA, which revealed a first principal component (PC1) that explained 39% of the variance in the data and split samples by species, followed by a second component (PC2) that explained 34% of the variance and primarily separated samples by brain region (Fig. 2b).

**Fig. 2.**
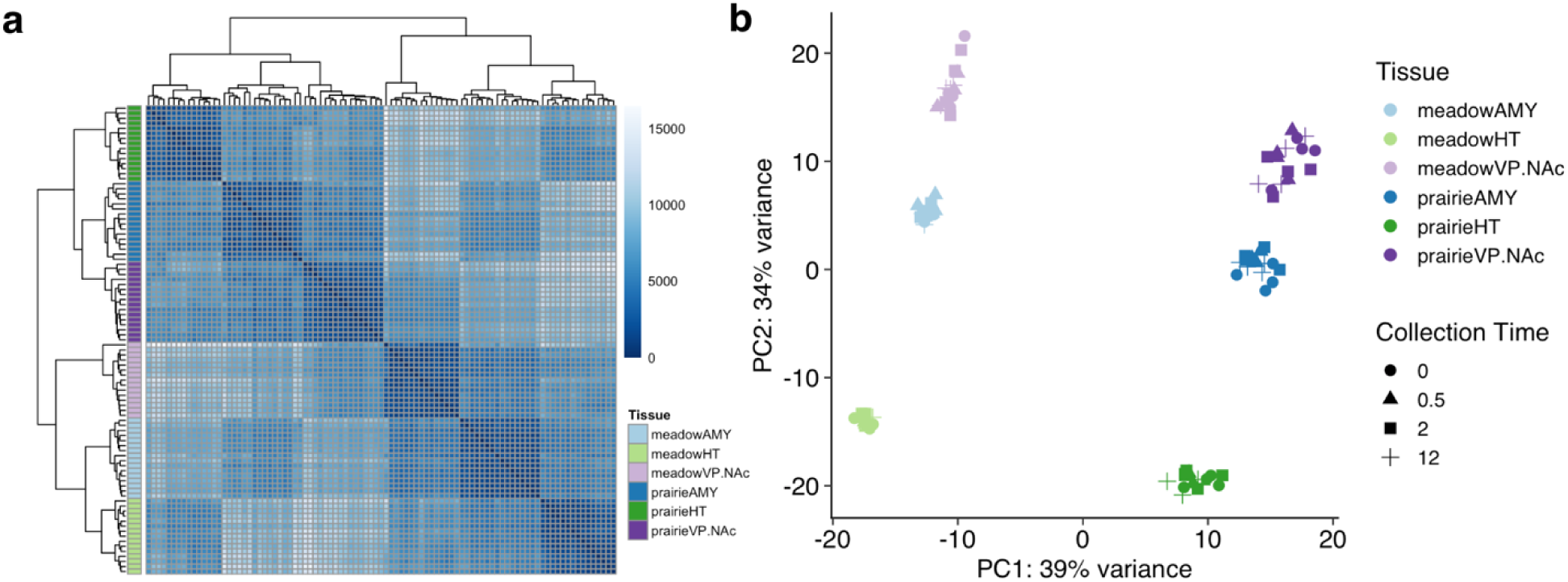
Vole brain gene expression varies consistently by species and region. **a.** Matrix of Poisson distances between each sample based on gene count data. Darker shades are more similar. Color key to the left shows sample type for each row. Hierarchical clustering trees show that samples split first by species, then by brain region. **b.** Principal component analysis of sample gene count data shows first principal component (PC1) that separates samples by species and accounts for 39% of sample variance. Second principal component (PC2) separates samples primarily by region and accounts for 34% of variance.

### Differential Gene Expression

#### Species Comparison of Virgins

To identify genes that may confer the capacity to bond in prairie but not meadow voles prior to mating, we used a subset of the data containing only samples collected from virgins (AMY, HT, and VP/NAc samples from n=4 individuals of each species). We tested for the effect of species using the likelihood ratio test implemented by DESeq2 (34). This test compares two generalized linear models fit to the data, a full model including all terms and a reduced model with one or more terms omitted. The test determines if the effect of the terms omitted in the reduced model is significantly greater than zero.

To determine the effect of species, we applied the full model ∼region + species and reduced model ∼region. 8377 of 12,672 genes were significantly differentially expressed (FDR<0.1) in this comparison (Fig. 3a, Supplementary Table 1). In order to identify the types of genes that differed in expression across species, we used the Mann-Whitney U (MWU) test (see Methods for details) to identify enriched gene ontology (GO) categories (35, 36). The MWU test of GO term enrichment revealed 48 significantly enriched (FDR<0.1) terms in the contrast between prairie and meadow vole virgins (Fig. 3b). This included 10 terms in the category Molecular Function, 11 in Biological Process, and 27 in the category Cellular Component. Enriched terms in prairie voles were largely associated with cell structure and extracellular space, while those enriched in meadow voles were associated with the ribosome and translation, mitochondrial function, and metabolic and biosynthetic processes.

**Fig. 3.**
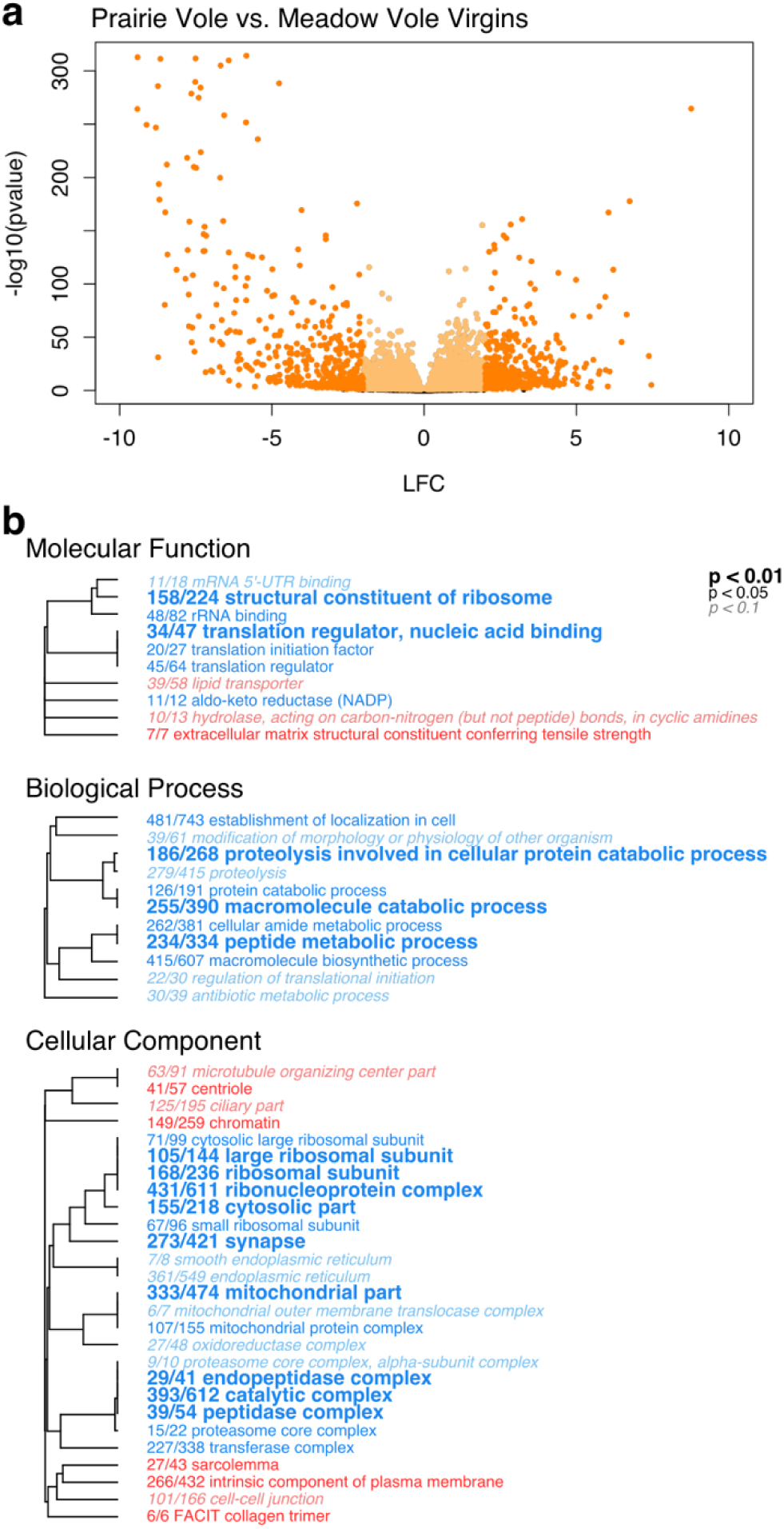
Vole brain gene expression varies by species. **a.** Volcano plot showing significance (-log_10_(p-value)) and magnitude of difference (log_2_ fold change, LFC) in expression for each gene in contrast between virgin prairie and meadow voles. Each point represents a single gene. Orange colored points pass significance threshold of FDR<0.1. Darker shaded points pass FDR cutoff and have LFC>2. Positive LFC values represent genes more highly expressed in prairie voles, negative LFC values are genes more highly expressed in meadow voles. **b.** Enriched GO terms for contrasts between prairie and meadow voles. Hierarchical clustering tree shows relationship between GO categories based on shared genes. Branches with length of zero are subsets of one another. Fractions preceding GO terms indicate proportion of “good” genes that have raw p-value<0.05 compared to total number of genes in the category. Bold text indicates adjusted p<0.01, plain text indicates adjusted p<0.05, and italicized text indicates adjusted p<0.1 for term. P-values are corrected using Benjamini-Hochberg false discovery rate procedure (85). Red terms are enriched in prairie voles and blue terms are enriched in meadow voles.

#### Comparison of Regions

As sample region was the second largest factor explaining gene expression data, we next compared expression across brain regions. To understand how gene expression differed across our focal brain regions, we used the entire dataset and applied the likelihood ratio test with the full model ∼species + region + time and the reduced model ∼species + time. Overall, 10,891 of 12,687 total genes significantly differed (FDR<0.1) in expression on the basis of sample region (Fig. 4a-c, Supplementary Table 2).

**Fig. 4.**
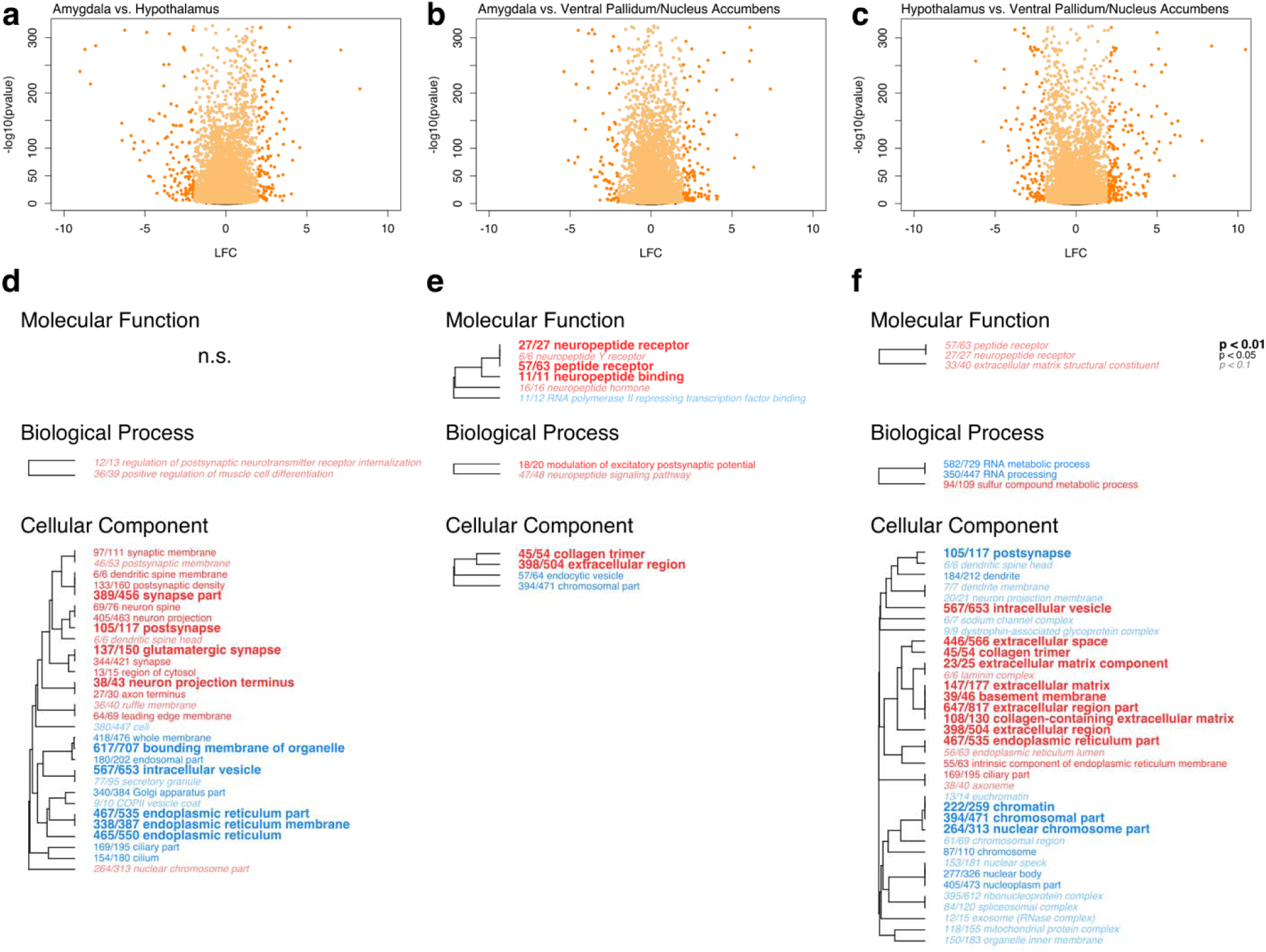
Vole brain gene expression varies by tissue type. **a-c.** Volcano plots showing significance (-log_10_(p-value)) and magnitude of difference (log_2_ fold change, LFC) in expression for each gene. Each point represents a single gene. Orange colored points pass significance threshold of FDR<0.1. Darker shaded points pass FDR cutoff and have LFC>2. **a.** Contrast between AMY and HT. Genes with positive LFC values are enriched in AMY, genes with negative LFC values are enriched in HT. **b.** Contrast between AMY and VP/NAc. Genes with positive LFC values are enriched in AMY, genes with negative LFC values are enriched in VP/NAc **c.** Contrast between HT and VP/NAc. Genes with positive LFC values are enriched in HT, genes with negative LFC values are enriched in VP/NAc. **d-f.** Enriched GO terms. Hierarchical clustering tree shows relationship between GO categories based on shared genes. Branches with length of zero are subsets of one another. Fractions preceding GO terms indicate proportion of “good” genes that have raw p-value<0.05 compared to total number of genes in the category. P-values are corrected using Benjamini-Hochberg false discovery rate procedure (85). Bold text indicates adjusted p<0.01, plain text indicates adjusted p<0.05, and italicized text indicates adjusted p<0.1 for term. n.s. indicates no significant terms in the category. **d.** Contrast between AMY and HT. Red terms enriched in AMY, blue terms enriched in HT. **e.** Contrast between AMY and VP/NAc. Red terms enriched in AMY, blue terms enriched in VP/NAc. **f.** Contrast between HT and VP/NAc. Red terms enriched in HT, blue terms enriched in VP/NAc.

Thirty-two GO terms were significantly enriched (MWU FDR<0.1) in the contrast between AMY and HT, including two in the category Biological Process and 30 under Cellular Component (Fig. 4d). Terms enriched in AMY were largely associated with neuronal structure and the synapse, while those enriched in HT were associated with membranous structures including endoplasmic reticulum, Golgi apparatus, intracellular vesicle, and whole membrane. Twelve GO terms were enriched in the contrast between AMY and VP/NAc (Fig. 4e). These included six in the category Molecular Function, two under Biological Process, and four in Cellular Component. Categories enriched in AMY were largely associated with neuropeptide signaling, including terms related to neuropeptide hormones and receptors, as well as extracellular space. Terms enriched in VP/NAc include *RNA polymerase II repressing transcription factor binding*, *endocytotic vesicle*, and *chromosomal part*. Finally, 42 terms were significantly enriched in the contrast between HT and VP/NAc, including three under Molecular Function, three in the category Biological Process, and 36 in the category Cellular Component (Fig. 4f). GO terms enriched in HT were largely associated with neuropeptide signaling and extracellular space, while those enriched in VP/NAc were related to synaptic structures as well as the nucleus, transcription, and translation.

#### Species Comparison Across Time

We next sought to determine the influence of mating and, in prairie voles, pair-bond formation on gene expression in our focal regions. In order to identify how gene expression changes differently across species following mating, we divided the full dataset into subsets containing all samples from a single region. We then tested these regional datasets with the likelihood ratio test, applying the full model ∼species + time + species:time and the reduced model ∼species + time. This comparison allowed us to identify those genes that exhibit species-specific patterns of expression following mating.

In the AMY, we found four genes that were significantly differentially expressed (FDR<0.1) across species over mating time points (Fig. 5a, Table 1, Supplementary Table 3). The gene with the smallest adjusted p-value (FDR=6.17*10^-5^) in this model was *Npas4*, an activity-dependent transcription factor involved in synapse formation (37) (Fig. 5d). In addition, *Nfkbia*, which encodes NF-Kappa-B Inhibitor Alpha, was significant in our models of both AMY and VP/NAc (Fig. 6a). We found 33 significantly enriched GO terms between prairie and meadow voles (MWU FDR<0.1), including two under Molecular Function and 31 in the category Cellular Component (Fig. 5g). Enriched terms in prairie voles were related to peptide receptors and the plasma membrane. Enriched terms in meadow voles were associated with ribosomal and mitochondrial function.

**Fig. 5.**
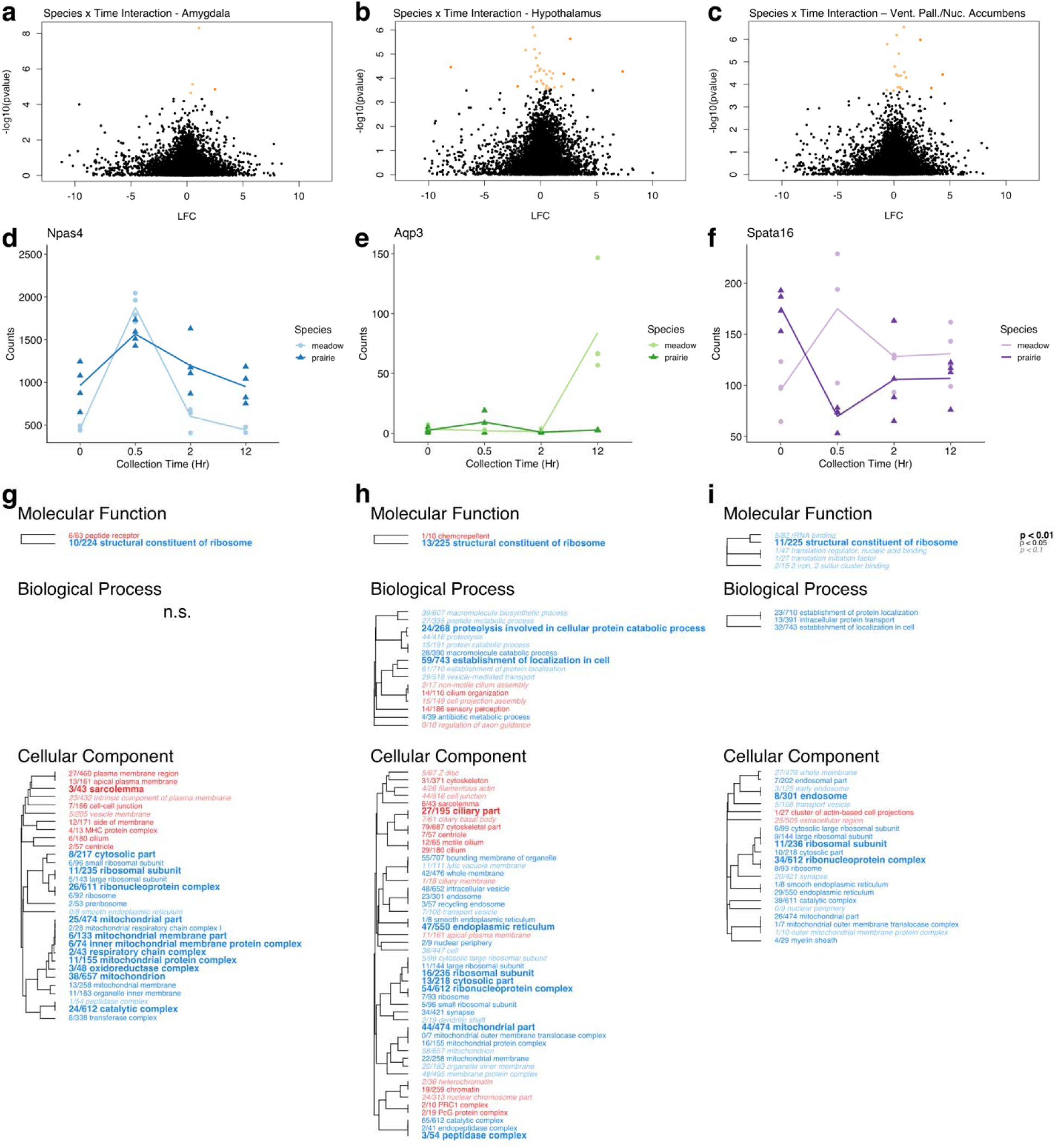
Vole brain gene expression varies over time by species. **a-c.** Volcano plots showing significance (-log_10_(p-value)) and magnitude of difference (log_2_ fold change, LFC) in expression for each gene in species:time interaction in **a.** AMY, **b.** HT, and **c.** VP/NAc. Each point represents a single gene. Orange colored points pass significance threshold of FDR<0.1. Darker shaded points pass FDR cutoff and have LFC>2. Genes with positive LFC values are enriched in prairie voles, genes with negative LFC values are enriched in meadow voles. **d-f.** Pattern of expression over time for genes with lowest adjusted p-value (FDR) in each regional model. Plots show counts for samples collected at each pre-mating (0 hr) and post-mating (0.5, 2, or 12 hr) collection point. Each point represents one sample. Darker shades and triangles represent prairie vole samples. Lighter shades and circles represent meadow vole samples. **d.** *Npas4* in AMY samples. **e.** *Aqp3* in HT samples. **f.** *Spata16* in VP/NAc samples. **g-i.** Enriched GO terms for species:time interaction in **g.** AMY, **h.** HT, and **i.** VP/NAc. Hierarchical clustering tree shows relationship between GO categories based on shared genes. Branches with length of zero are subsets of one another. Fractions preceding GO terms indicate proportion of “good” genes that have raw p-value<0.05 compared to total number of genes in the category. P-values are corrected using Benjamini-Hochberg false discovery rate procedure (85). Red terms enriched in prairie voles, blue terms enriched in meadow voles. Bold text indicates adjusted p<0.01, plain text indicates adjusted p<0.05, and italicized text indicates adjusted p<0.1 for term. n.s. indicates no significant terms in the category.

**Fig. 6.**
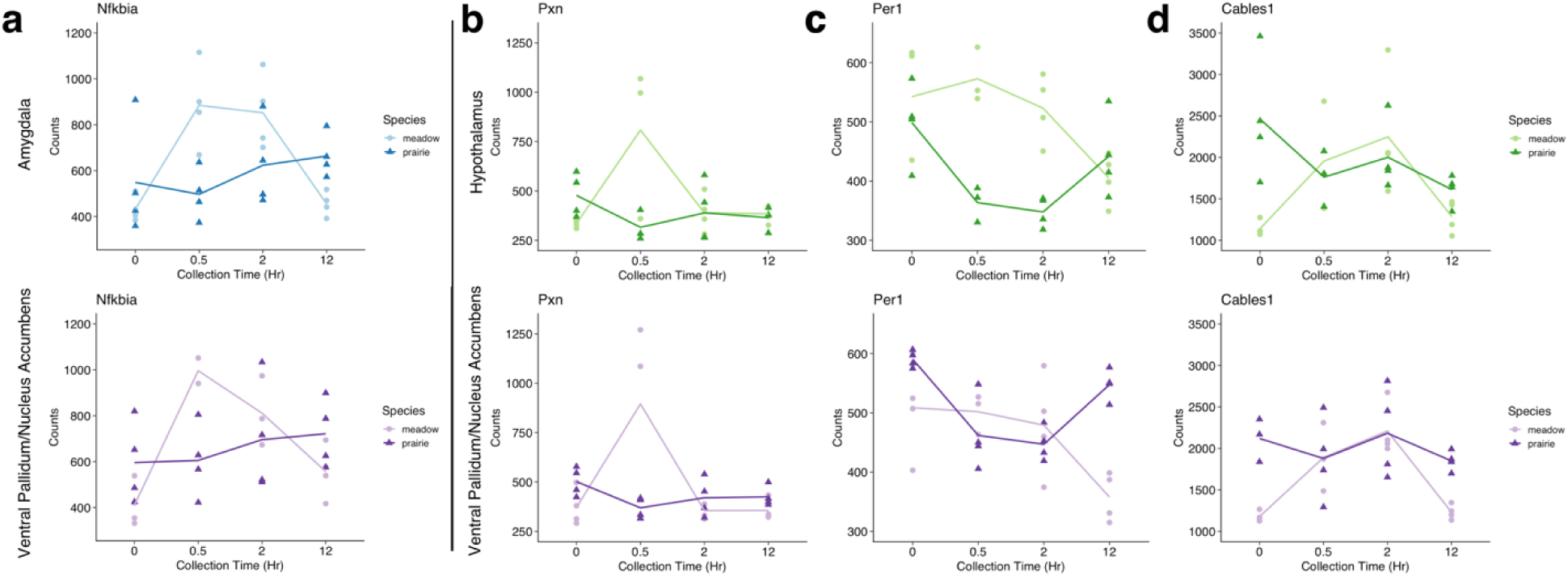
Expression pattern of significant genes is consistent across regions. Plots show counts for samples collected at each pre-mating (0 hr) and post-mating (0.5, 2, or 12 hr) collection point. Each point represents one sample. Darker shades and triangles represent prairie vole samples. Lighter shades and circles represent meadow vole samples. **a.** *Nfkbia* in AMY (above) and VP/NAc (below). **b.** *Pxn* in HT (above) and VP/NAc (below). **c.** *Per1* in HT (above) and VP/NAc (below). **d.** *Cables1* in HT (above) and VP/NAc (below).

**Table 1.**
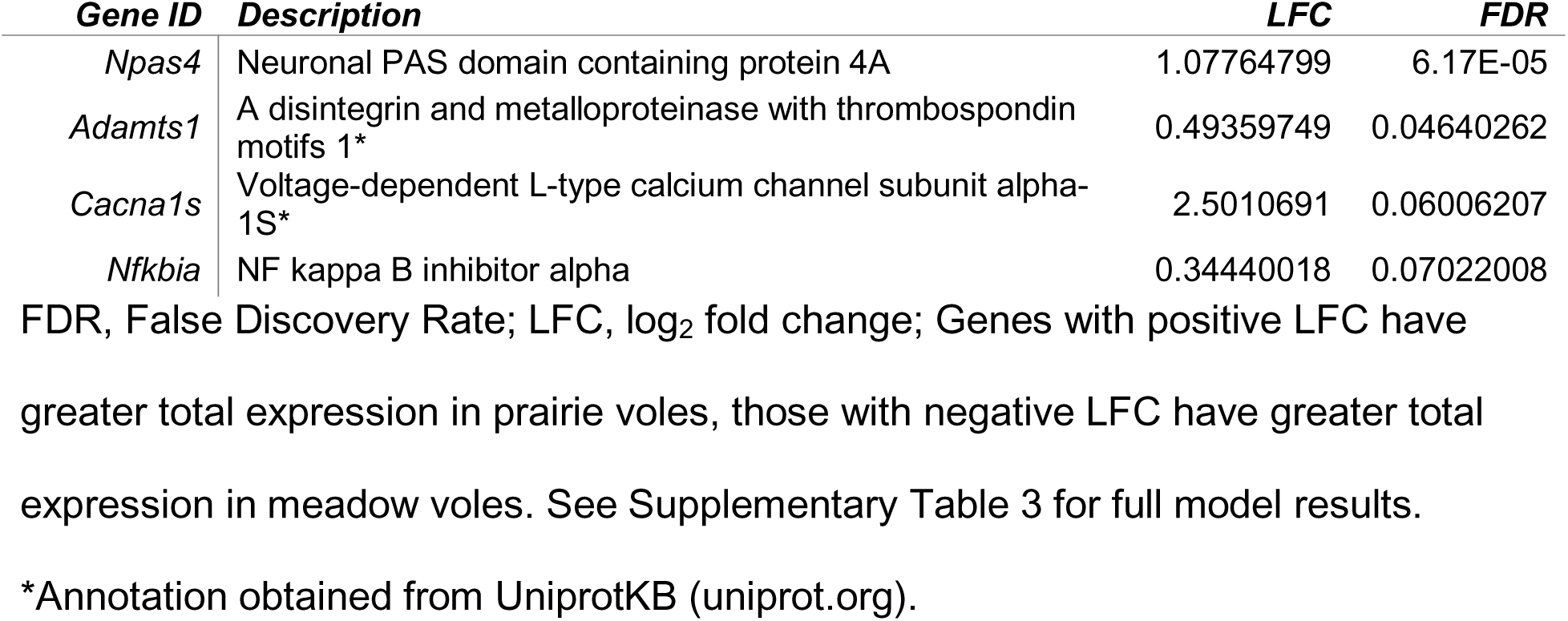
Differentially expressed genes in amygdala over time across species

| *Gene ID* | *Description* | *LFC* | *FDR* |
| --- | --- | --- | --- |
| *Npas4* | Neuronal PAS domain containing protein 4A | 1.07764799 | 6.17E-05 |
| *Adamts1* | A disintegrin and metalloproteinase with thrombospondin motifs 1* | 0.49359749 | 0.04640262 |
| *Cacna1s* | Voltage-dependent L-type calcium channel subunit alpha-1S* | 2.5010691 | 0.06006207 |
| *Nfkbia* | NF kappa B inhibitor alpha | 0.34440018 | 0.07022008 |

FDR, False Discovery Rate; LFC, log<sub>2</sub> fold change; Genes with positive LFC have greater total expression in prairie voles, those with negative LFC have greater total expression in meadow voles. See Supplementary Table 3 for full model results.
\*Annotation obtained from UniprotKB (uniprot.org).

In the HT, 31 genes were significantly differentially expressed (FDR<0.1) across species over mating time points (Fig. 5b, Supplementary Table 4), of which 25 were annotated (Table 2). These included water channel-encoding gene *Aqp3* (38) (Fig. 5e), which had the lowest adjusted p-value (FDR=0.0089713), as well as *Pxn*, which encodes paxillin, an adaptor protein that interacts with the cytoskeleton and signaling molecules to facilitate neurite outgrowth and long-term potentiation (39, 40) (Fig. 6b), the circadian clock gene *Per1* (41) (Fig. 6c), and *Cables1* which encodes a kinase-binding protein that promotes neurite outgrowth (42) (Fig. 6d), each of which were significant in the VP/NAc model as well. 65 GO terms were significantly enriched (MWU FDR<0.1) in the contrast between prairie and meadow voles. Among these were two in the category Molecular Function, 15 in Biological Process, and 48 in Cellular Component (Fig. 5i). Enriched terms in prairie voles were typically associated with neuronal development and cell structure. Terms enriched in meadow voles included many related to ribosomal and mitochondrial function, metabolic and biosynthetic processes, and vesicular transport.

**Table 2.**
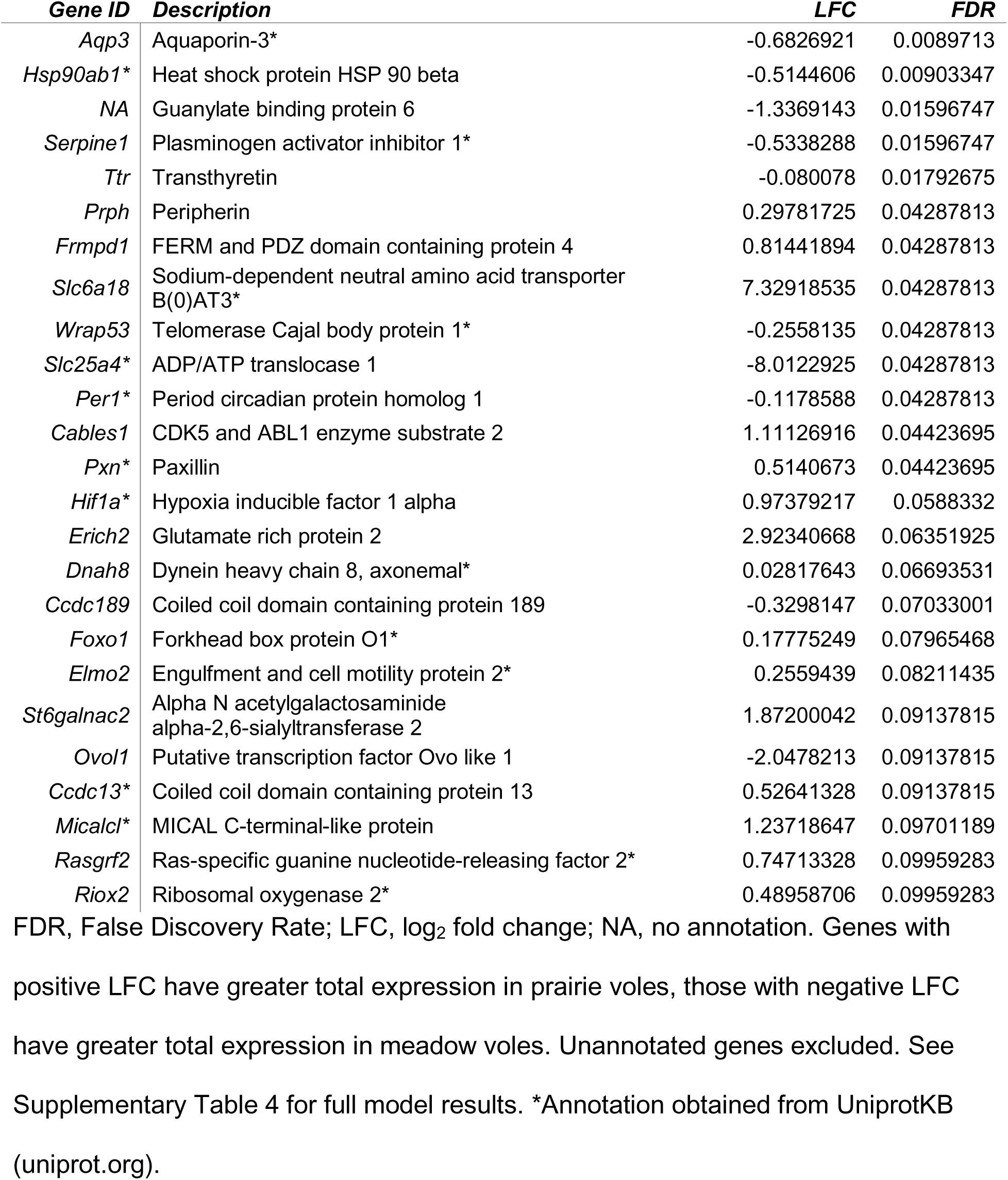
Differentially expressed genes in hypothalamus over time across species

In the VP/NAc we found 21 genes that were significantly differentially expressed over mating time points (Fig. 5c, Supplementary Table 5), 19 of which were annotated (Table 3). Among these, *Spata16*, a gene with known function in spermatogenesis (43) (Fig. 5f) had the lowest adjusted p-value (FDR=0.00318981). Several genes that were significant in the VP/NAc model were also significant in models for other regions. These include *Nfkbia* (Fig. 6a), which was also differentially expressed in AMY, as well as *Pxn* (Fig. 6b), *Per1* (Fig. 6c), and *Cables1* (Fig. 6d), which were each also differentially expressed in HT. In addition, we found 30 significantly enriched (MWU FDR<0.1) GO terms between prairie and meadow voles, which included five under Molecular Function, three under Biological Process, and 22 in Cellular Component (fig. 5i). Only two of these terms were enriched in prairie voles: *cluster of actin-based cell projections* and *extracellular region*. Many of the terms enriched in meadow voles were associated with translation, protein localization, mitochondria, and the endosome.

**Table 3.** Differentially expressed genes in ventral pallidum/nucleus accumbens over time across species

| *Gene ID* | *Description* | *LFC* | *FDR* |
| --- | --- | --- | --- |
| *Spata16* | Spermatogenesis-associated protein 16* | 0.88879921 | 0.00318981 |
| *Dhx35* | Probable ATP dependent RNA helicase DHX35 | 0.2499954 | 0.00366884 |
| *Mccc2* | Methylcrotonoyl-CoA carboxylase beta chain, mitochondrial* | -0.6041633 | 0.00462278 |
| *Pxn\** | Paxillin | 0.43629243 | 0.00956998 |
| *Sike1* | Suppressor of IKBKE 1 | 0.06660377 | 0.03030073 |
| *Rbm3* | RNA binding protein 3 | 0.38113582 | 0.03897143 |
| *P4ha2* | Prolyl 4-hydroxylase subunit alpha-2* | 0.93405983 | 0.03897143 |
| *Best3* | Bestrophin 2 | 4.3486464 | 0.03897143 |
| *Per1\** | Period circadian protein homolog 1 | 0.21428454 | 0.03897143 |
| *Ciart* | Circadian associated transcriptional repressor | 0.31693109 | 0.03897143 |
| *Agt* | Angiotensinogen* | 0.61071671 | 0.03897143 |
| *Prss56* | Transmembrane protease serine 3 | 1.07993336 | 0.04014217 |
| *Cables1* | CDK5 and ABL1 enzyme substrate 2 | 1.09018372 | 0.04014217 |
| *Sgk1* | Serine/threonine-protein kinase Sgk1* | 0.34324947 | 0.09405603 |
| *Nfkbia* | NF kappa B inhibitor alpha | 0.53419379 | 0.09405603 |
| *Adamts14* | NA | 3.33669686 | 0.09496818 |
| *Csrnp1* | Cysteine/serine rich nuclear protein 3 | 0.59391793 | 0.09496818 |
| *Lrrc6* | Protein tilB homolog* | 0.71670559 | 0.09496818 |
| *Smoc1* | SPARC related modular calcium binding protein 2 | -0.6408534 | 0.09995594 |

FDR, False Discovery Rate; LFC, log<sub>2</sub> fold change; NA, no annotation. Genes with positive LFC have greater total expression in prairie voles, those with negative LFC have greater total expression in meadow voles. Unannotated genes excluded. See Supplementary Table 5 for full model results. \*Annotation obtained from UniprotKB (uniprot.org).

### Correlated Gene Networks

We next sought to determine if co-regulated groups of genes varied in expression related to our variables of interest (sample region, mating status [virgin vs. mated], or time of collection after mating onset). Using the R package WGCNA (44), we constructed gene expression networks for prairie and meadow voles, and detected modules of genes with correlated expression patterns (Fig. 7, see Tables 4 and 5 for Pearson r and p-values for significant correlations). For prairie voles, we detected 16 modules ranging in size from 72 to 1707 genes as well as 392 that were unassigned to any module. In meadow voles we detected 12 modules ranging in size from 167 to 1577 genes and 517 genes that were unassigned. We then correlated these modules to sample traits including brain region (AMY, HYP, or VP/NAc), mating status (mated vs. virgin), and collection time points, including time as a continuous variable and each post-mating time point individually.

**Fig. 7.**
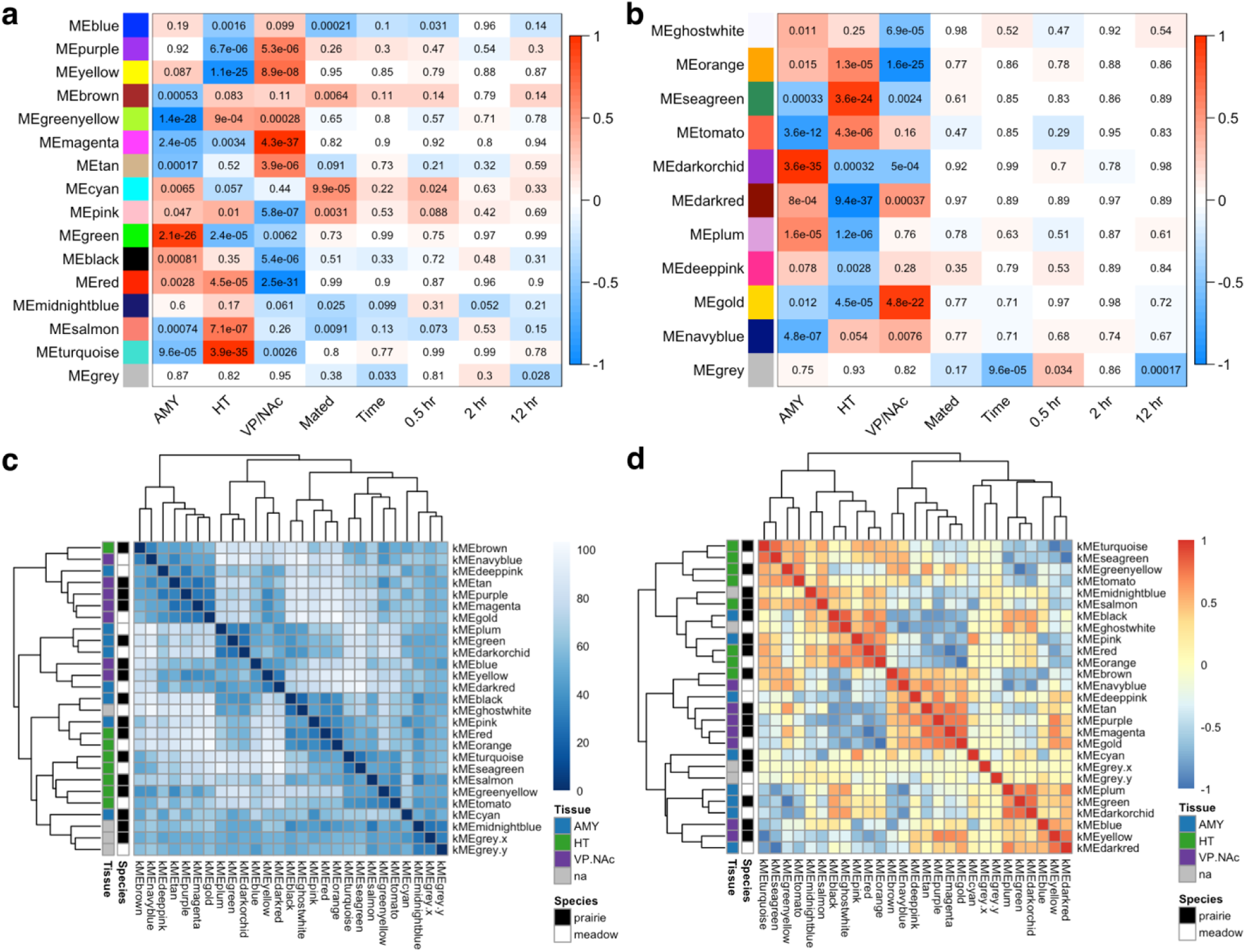
Correlated gene networks form modules related to tissue type and mating status. **a-b.** Correlations between WGCNA modules and sample traits including tissue (AMY, HT, or VP/NAc), mating status (Mated), collection time point as a continuous variable (Time) or individual post-mating time points (0.5 hr, 2 hr, or 12 hr) for **a.** prairie voles and **b.** meadow voles. Color scale bar indicates Pearson r value for correlation. Red indicates positive r value and blue indicates negative r value. Each cell in matrix shows correlation p-value. See Tables 4 & 5 for Pearson r values for significant correlations. **c.** Matrix of distances between sample module membership scores (kME). Color bar indicates sample distances with darker shades indicating more similar samples. Dendrograms show hierarchical clustering of samples based on distance. **d.** Matrix of correlations between sample module membership scores (kME). Color bar indicates sample correlations with reds indicating positively correlated samples and blues indicating negatively correlated samples. Dendrograms show hierarchical clustering of samples based on distance. Color legends in **c** and **d** indicate module species (black=prairie vole, white=meadow vole) and the brain region most positively correlated with the module (blue=AMY, green=HT, purple=VP/NAc, grey=no significant positive correlation with brain region)

**Table 4.**
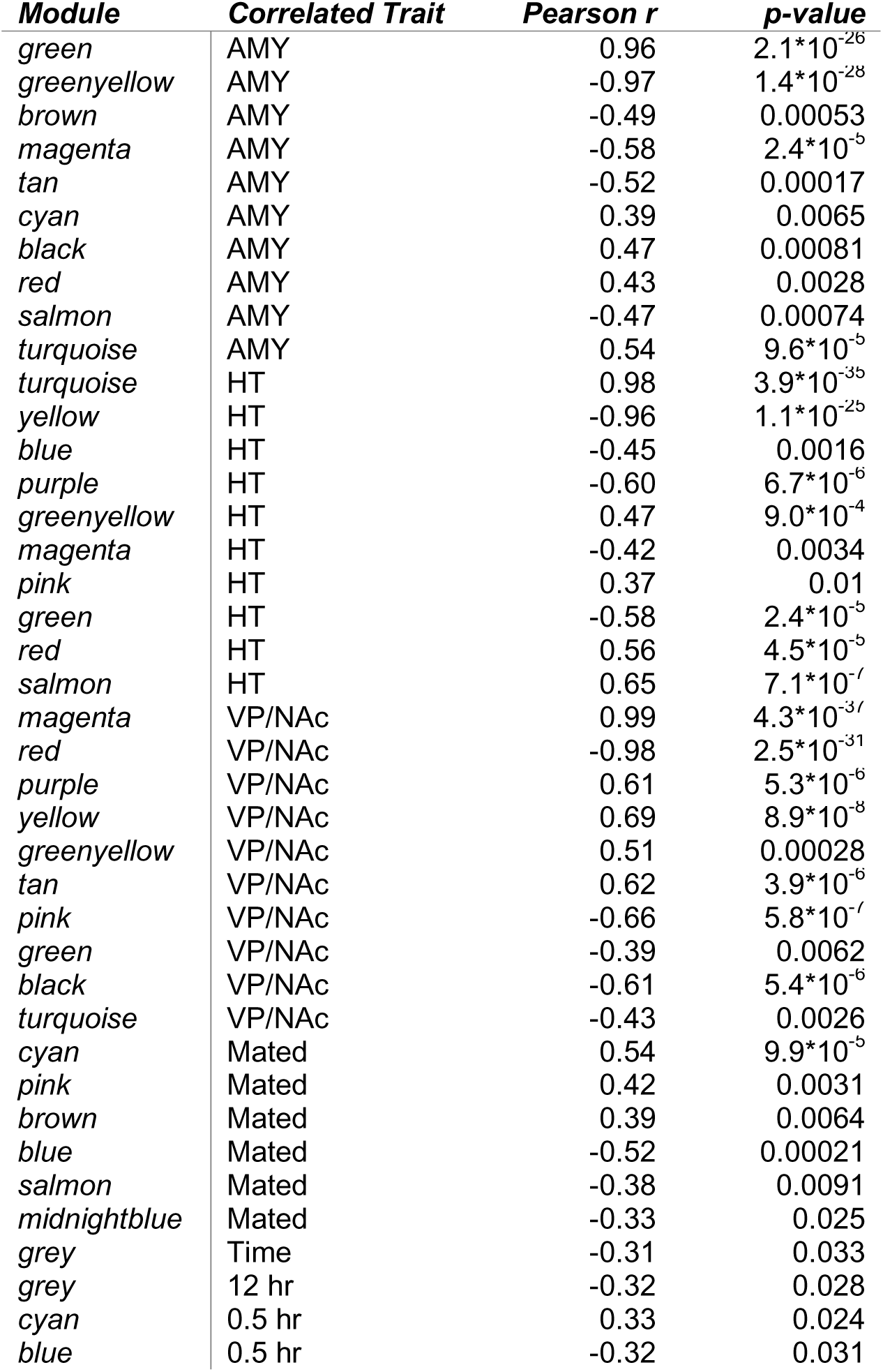
Statistics for prairie vole WGCNA modules with significant correlations

**Table 5.**
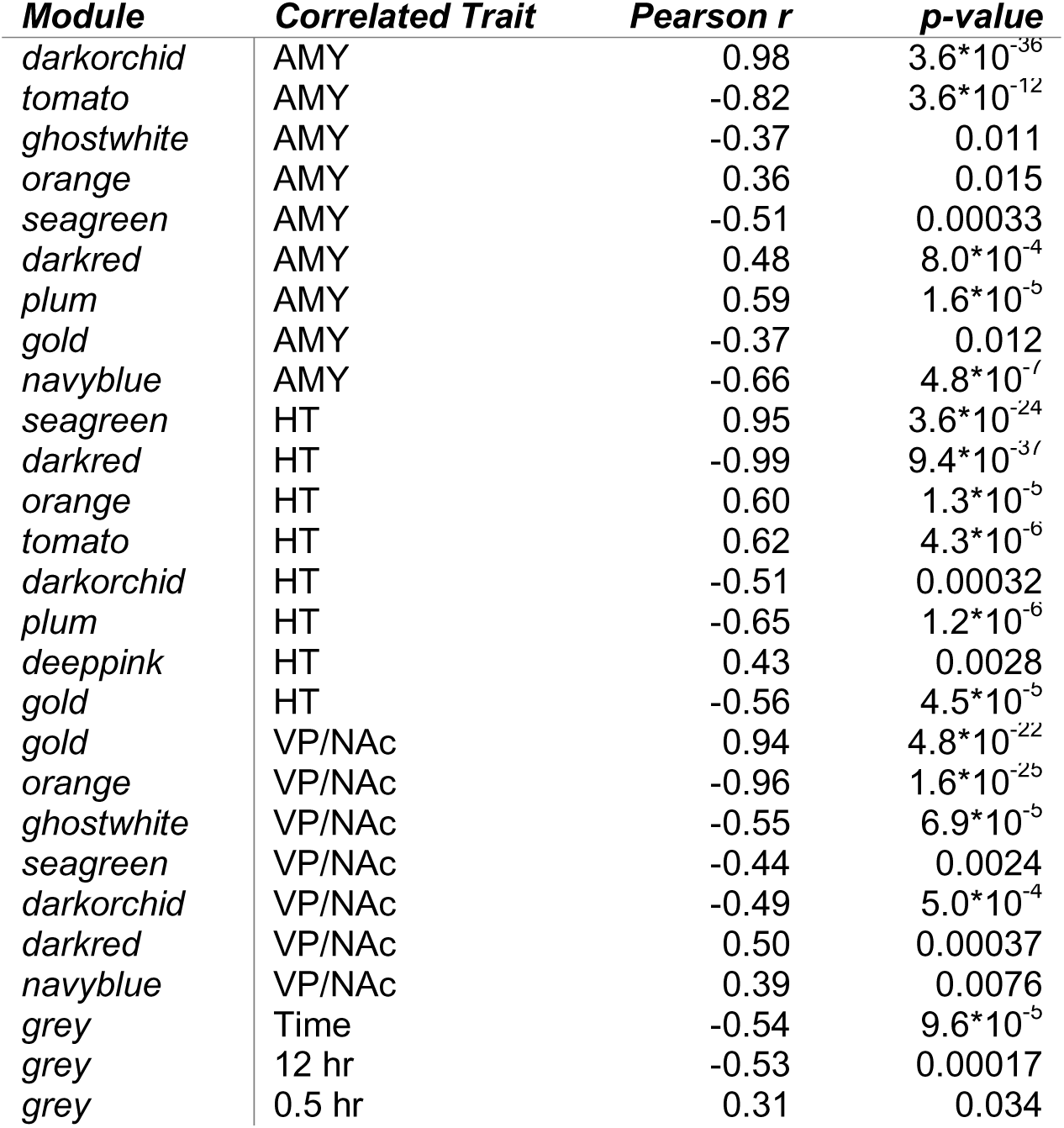
Statistics for meadow vole WGCNA modules with significant correlations

#### Module correlation with sample traits

For both species, the traits most strongly correlated with modules were sample brain region (Fig. 7a, b). In prairie voles (Table 4), the green module was most strongly positively correlated with AMY, while the greenyellow module was strongly negatively correlated with the region. The brown, magenta, tan, cyan, black, red, salmon, and turquoise modules were also significantly correlated with AMY. For HT, the turquoise module showed the strongest positive correlation, and yellow had the strongest negative correlation. Modules blue, purple, greenyellow, magenta, pink, green, red, and salmon were also significantly correlated with HT. Finally, the magenta module was most strongly positively, and red module most strongly negatively correlated with VP/NAc. The purple, yellow, greenyellow, tan, pink, green, black, and turquoise modules were also significantly correlated with this region.

In meadow voles (Table 5), the darkorchid module was most strongly positively, and tomato most strongly negatively correlated with AMY. The ghostwhite, orange, seagreen, darkred, plum, gold, and navyblue modules were also significantly correlated with AMY. Modules seagreen and darkred were most positively and negatively correlated with HT, respectively. The orange, tomato, darkorchid, plum, deeppink, and gold modules also had significant correlations with HT. Finally, the gold module had the strongest positive correlation with VP/NAc and orange module had the strongest negative correlation. Modules ghostwhite, seagreen, darkorchid, darkred, and navyblue each also significantly correlated with VP/NAc.

In general, module correlations with mating status and collection time points were weaker than those with brain regions (Fig. 7a, b). For prairie voles (Fig. 7a, Table 4) three modules, cyan, pink, and brown were significantly positively correlated with having mated. The blue, salmon, and midnightblue modules were negatively correlated with having mated, indicating a positive association with virgin animals. Only the grey module containing unassigned genes was significantly correlated with overall time of collection in the experiment. This relationship appears to be driven by a significant negative correlation with the 12 hr collection time point. However, two modules were significantly correlated with specific collection time points. The cyan module was positively associated with the 0.5 hr time point and the blue module was significantly negatively correlated with this time.

Module correlations with mating status and post-mating time points were weaker in meadow voles than in prairie voles (Fig. 7b, Table 5). No modules were significantly correlated with mating status, and only the grey module containing unassigned genes was correlated with collection time or individual time points. It was negatively correlated with time overall and the 12 hr collection time point, and positively correlated with the 0.5 hr collection time point.

#### Module associations across species

To identify relationships between modules across species, we clustered prairie and meadow vole modules on the basis of Euclidean distance (Fig. 7c) and Pearson correlation (Fig. 7d). Clustering revealed eight pairs of modules that were closely related across species. These included the brown prairie vole and navyblue meadow vole modules, prairie magenta and meadow gold, prairie green and meadow darkorchid, prairie yellow and meadow darkred, prairie black and meadow ghostwhite, prairie red and meadow orange, prairie turquoise and meadow seagreen, and prairie greenyellow and meadow tomato. Clustering by correlation also found one pair of prairie vole modules, midnightblue and salmon, that were most closely related.

Of these closely related modules, the majority were strongly associated with individual brain regions. The green/darkorchid modules were the most strongly positively correlated with AMY in both species, while the greenyellow/tomato modules were the most negatively correlated with AMY. Modules turquoise/seagreen were most highly positively correlated with HT and yellow/darkred were most strongly negatively correlated with it. Finally, the gold/magenta modules were most strongly associated and red/orange modules most strongly negatively associated with VP/NAc.

We used GO term analysis on pairs of modules that were strongly positively correlated with each region to identify enriched gene categories. The prairie vole green module, which was strongly correlated with AMY had 85 significantly enriched GO terms (MWU FDR<0.1), including four in the category Molecular Function, 42 under Biological Process, and 39 in the category Cellular Component (Fig. 8a). In meadow voles, the AMY-associated darkorchid module had 53 significantly enriched (MWU FDR<0.1) GO terms (Fig. 8b.). These included one term in Molecular Function, *cation channel activity*, 27 terms related to Biological Process, and 25 terms in the category Cellular Component. In both species, terms related to neuronal projections, ion channels, plasma membrane and extracellular space, receptors, and synaptic transmission were enriched. The terms *behavior* and *cognition* were also enriched in both the prairie vole green module and meadow vole darkorchid module. However, the terms *associative learning* and *learning* were uniquely enriched in the prairie vole green module.

**Fig. 8.**
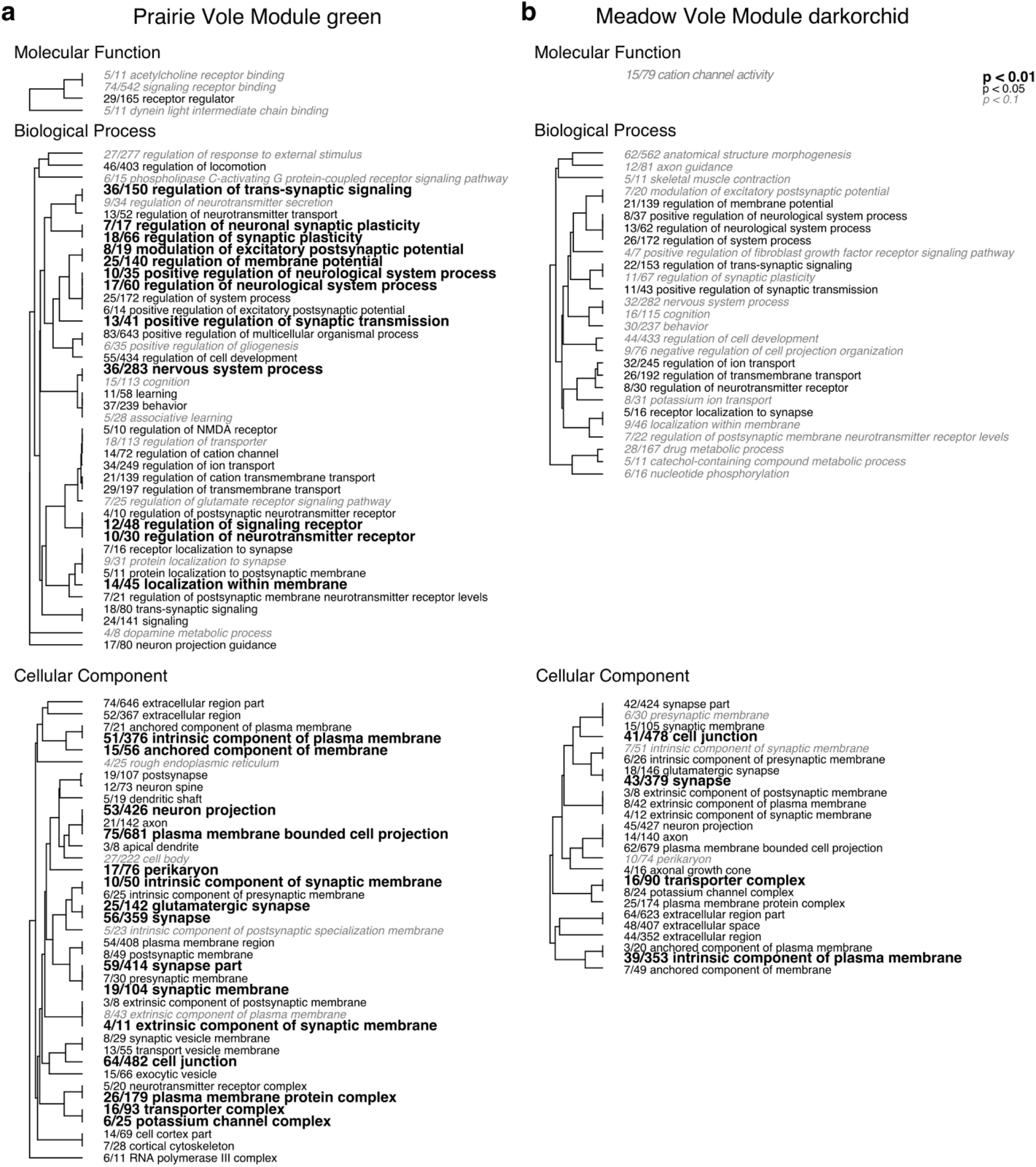
Gene ontology analysis for amygdala gene-expression modules. Enriched GO terms in **a.** prairie vole green module and **b.** meadow vole darkorchid module. Hierarchical clustering tree shows relationship between GO categories based on shared genes. Branches with length of zero are subsets of one another. Fractions preceding GO terms indicate proportion of genes from the category that are included in the module of interest. FDR determined by 100 permutations where significance measures are randomly shuffled among genes. Bold text indicates adjusted p<0.01, plain text indicates adjusted p<0.05, and italicized text indicates adjusted p<0.1 for term. n.s. indicates no significant terms in the category.

The prairie vole module turquoise and meadow vole module seagreen were each strongly associated with HT. 33 GO terms were significantly enriched in the turquoise module (MWU FDR<0.1), including 12 in the category Molecular Function, nine in Biological Process, and 12 in Cellular Component (Fig. 9a). In the meadow vole seagreen module, 11 terms were enriched (MWU FDR<0.1), all under Cellular Component (Fig. 9b). In both species, several terms related to cilia were enriched. Additionally, in prairie voles, terms relating to neuropeptide signaling, transcription factors, extracellular space were enriched along with the term *feeding behavior*.

**Fig. 9.**
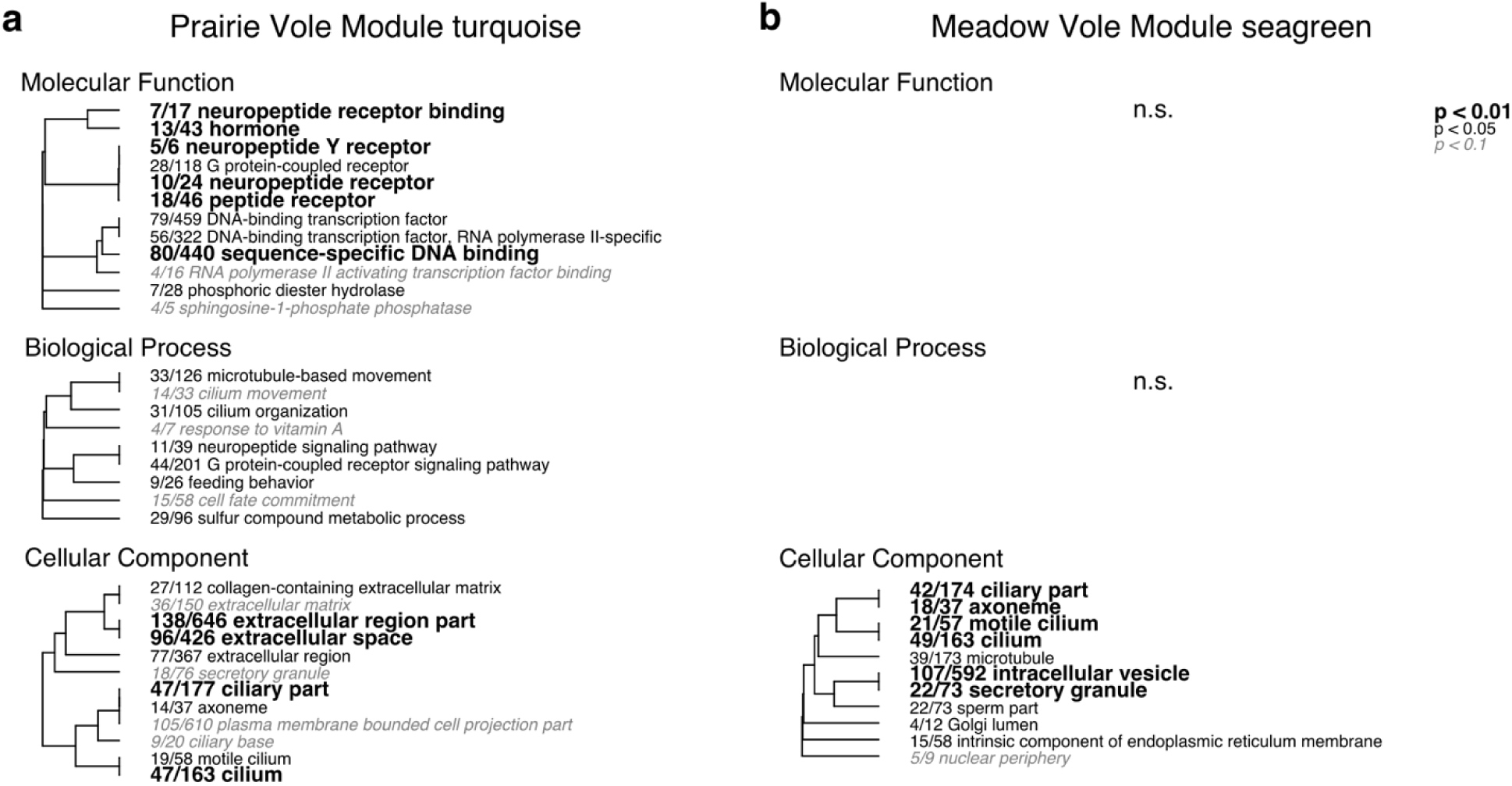
Gene ontology analysis for hypothalamic gene-expression modules. Enriched GO terms in **a.** prairie vole tuquoise module and **b.** meadow vole seagreen module. Hierarchical clustering tree shows relationship between GO categories based on shared genes. Branches with length of zero are subsets of one another. Fractions preceding GO terms indicate proportion of genes from the category that are included in the module of interest. FDR determined by 100 permutations where significance measures are randomly shuffled among genes. Bold text indicates adjusted p<0.01, plain text indicates adjusted p<0.05, and italicized text indicates adjusted p<0.1 for term. n.s. indicates no significant terms in the category.

No GO terms were significantly enriched in the prairie vole magenta module, which was strongly correlated with VP/NAc. However, 12 terms were significantly enriched (MWU FDR<0.1) in the related meadow vole gold module (Fig. 10). These included five in the category Molecular Function and six in the category Cellular Component. In addition, one term, *regulation of postsynaptic neurotransmitter receptor internalization* was enriched under the category Biological Process. Overall, most enriched terms related to the cell membrane, synaptic transmission, or enzymatic function.

**Fig. 10.**
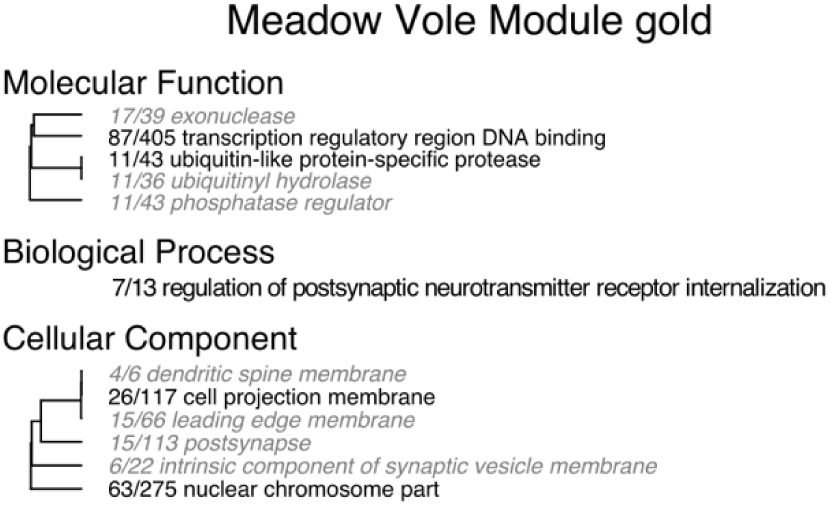
Gene ontology analysis for ventral pallidum/nucleus accumbens gene-expression modules. Enriched GO terms in meadow vole gold module. (There were no significant enrichments for the prairie vole magenta module.) Hierarchical clustering tree shows relationship between GO categories based on shared genes. Branches with length of zero are subsets of one another. Fractions preceding GO terms indicate proportion of genes from the category that are included in the module of interest. FDR determined by 100 permutations where significance measures are randomly shuffled among genes. Bold text indicates adjusted p<0.01, plain text indicates adjusted p<0.05, and italicized text indicates adjusted p<0.1 for term. n.s. indicates no significant terms in the category.

#### Modules associated with mating

Finally, to identify gene categories associated with mating, we looked for GO term enrichment among the modules most strongly correlated with mating status in prairie voles (Fig. 11). We used the MWU test to identify significantly enriched terms in the cyan and pink modules, which were the most significantly positively correlated with having mated, and the blue and salmon modules, which were the two most significantly negatively correlated with having mated (thus positively associated with virgin animals).

**Fig. 11.**
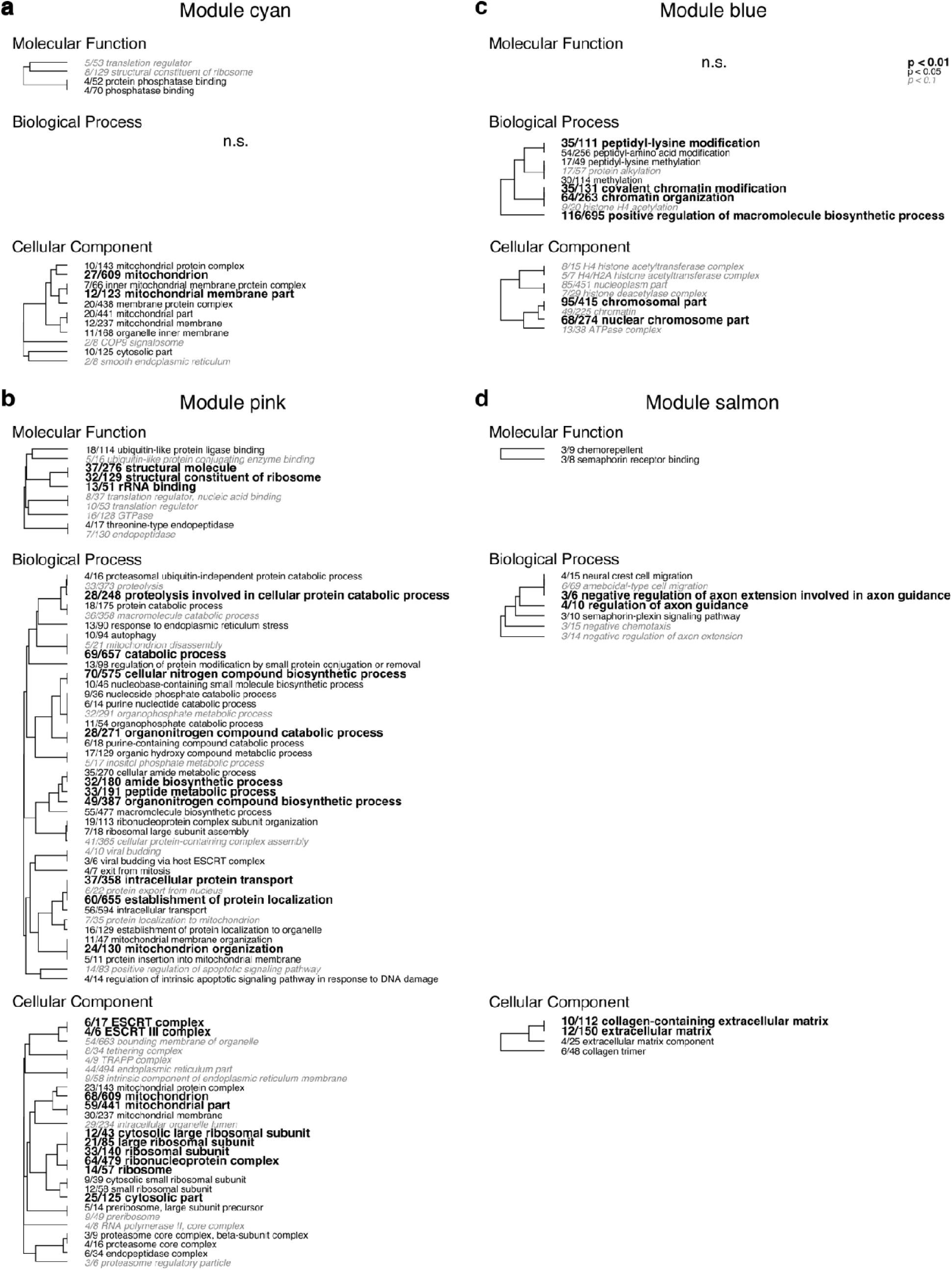
Effects of mating on gene ontology categories. Enriched GO terms in modules most strongly positively (**a-b**) and negatively (**c-d**) correlated with mating status in prairie voles. **a.** cyan module, **b.** pink module **c.** blue module, **d.** salmon module. Hierarchical clustering tree shows relationship between GO categories based on shared genes. Branches with length of zero are subsets of one another. Fractions preceding GO terms indicate proportion of genes from the category that are included in the module of interest. FDR determined by 100 permutations where significance measures are randomly shuffled among genes. Bold text indicates adjusted p<0.01, plain text indicates adjusted p<0.05, and italicized text indicates adjusted p<0.1 for term. n.s. indicates no significant terms in the category.

In addition to mating status, these modules also had significant correlations with brain regions (Fig. 7a, Table 4). The cyan module was positively correlated with AMY, pink module was positively correlated with AMY and HT, and negatively correlated with VP/NAc. Module blue was negatively correlated with HT and the salmon module was positively correlated with HT and negatively correlated with AMY.

Both the cyan and pink modules, each had significantly enriched terms (MWU FDR<0.5) related to translation, ribosomes, and mitochondria (Fig. 11a, b). In addition, terms associated with phosphatase binding were enriched in the cyan module and several terms related to ubiquitination, catabolic and metabolic processes, protein localization, and the ESCRT complex were enriched in the pink module. Terms significantly enriched (MWU FDR<0.5) in the blue included several related to peptidyl-amino acid modification and chromatin modification (Fig. 11c). Those enriched in the salmon module (MWU FDR<0.5) were largely associated with neuronal development and axon guidance, as well as extracellular space (Fig. 11d).

## Discussion

Pair bonding is an important social behavior of many animal species, including humans (1,4,45,46). Significant efforts have been made to understand the neural and hormonal mechanisms underlying the formation of pair bonds; however, these studies have focused on a relatively small number of genes and their products (17). A critical next step in understanding the mechanisms of bond formation is more fully accounting for the genes underlying pair-bond formation. While other studies have made comparisons of neural gene expression in closely-related monogamous and non-monogamous species (47), to our knowledge, ours is the first to make comparisons across species during an actively-developing bond, as well as the first to focus on specific regions of interest, rather than the entire brain.

In this study, we used RNA-sequencing to observe changes in gene expression in three regions that have critical roles in pair-bond formation including social memory, social context, and reward. We compared two species, the bond-forming prairie vole and the non-bonding meadow vole, before and at three time points after mating. By using a comparative approach, we sought to differentiate those genes which change in expression in response to mating, but do not play a role in bond formation, from the genes that may be critical to pair-bond development. We first identified differences in the gene categories expressed across species in virgins, as well as differences in the categories of genes expressed across our focal regions. Next, we identified those genes that differed in expression post-mating across species in each focal region. Finally, we identified gene expression networks in each species and related modules of genes with correlated expression to sample features. This allowed us to identify modules strongly associated with each brain region, as well as modules which changed in expression in response to mating in prairie voles.

### Pre-mating differences and the capacity to bond

Our results emphasize the importance of pre-mating differences in gene expression that confer the capacity to bond in prairie voles but not meadow voles. We found that gene expression count data were most strongly associated with sample species (Fig 2), and the vast majority of genes included in this study were differentially expressed across prairie and meadow vole virgins (Fig 3). This is consistent with prior studies that have investigated the neural mechanisms of pair bonding through comparisons between monogamous and promiscuous vole species. For example, the neuropeptide AVP plays an important role in the development of partner preference and aggression toward unfamiliar conspecifics that characterize pair-bond formation in prairie voles (26). Expression of the V1a vasopressin receptor in the VP is necessary for bond formation in male voles (48). Meadow voles naturally have very low expression of this receptor in the VP, however, driving its expression in this brain region results in increased partner preference, a hallmark of bond formation, in meadow voles (12).

Similarly, several other studies focused on individual candidate genes or proteins found species differences in expression of OT receptor, mu-opioid receptor, corticotropin-releasing factor receptors, estrogen receptor alpha, and neuropeptide Y (49–53). Gene expression differences important for behavior have also been found across bonding and non-bonding species in other taxa, including variation in *Avp* (gene encoding arginine-vasopressin) expression related to parental care in *Peromyscus* mice (54) and variation in OT receptor expression across pair-bonding versus solitary species of butterflyfish (55). Together, these and the present study show that species-level differences in brain gene expression play a large role in determining the capacity to form pair bonds.

While the effects of vasopressin and its receptor are dramatic, it remains likely that many genes underlie the species differences in bonding capacity. Our current study has identified several gene categories that differed in expression across species in virgins. GO terms enriched in prairie voles were largely related to cell structure, the nucleus and extracellular space. These include *microtubule organizing center part*, *centriole, chromatin,* and *cell-cell junction.* Enriched in meadow voles were terms associated with the ribosome and translation, mitochondrial function, and metabolic and biosynthetic processes including *translation initiation factor*, *translation regulator*, *peptide metabolic process*, *macromolecule biosynthetic process*, *ribosomal subunit*, and *mitochondrial part*. These results indicate that gene expression differences across virgin prairie voles are largely associated with genes involved in cell structure (enriched in prairie voles) and protein synthesis (enriched in meadow voles).

### Function of critical bonding-related regions

While it was not the central focus of this study, given that we found sample brain region to be the second strongest factor associated with gene expression, we explored the types of genes that differed between our focal brain regions. Unsurprisingly, given its role in social memory (23), when comparing the AMY to HT, we found enrichment of terms related to synaptic signaling including *regulation of postsynaptic neurotransmitter receptor internalization*, *postsynapse*, and *glutamatergic synapse* (Fig. 4d). In comparison to the VP/NAc, many enriched GO terms in AMY were related to neuropeptide signaling (Fig. 4e). Neuropeptide signaling, particularly by OT, is necessary for the AMY’s function in social recognition and memory. The medial AMY expresses OT receptor and receives OT innervation from the paraventricular nucleus of the HT in both mice and prairie voles (23, 29). OT knockout mice are unable to recognize familiar conspecifics, but this deficit is rescued by site-specific injection of OT into the medial AMY (23). Further, site-specific injection of an OT antagonist into the medial AMY also disrupted social recognition in wild type mice. Beyond OT, the AMY expresses many other neuropeptides and their receptors (56). Genes in the neuropeptide-related categories enriched in AMY included those encoding galanin, neuropeptide FF, urocortin-3 along with receptors for neuropeptide Y, galanin, somatostatin, and neuropeptide FF along with several others. Our results suggest that there are a variety of neuropeptide systems that are likely at play in AMY function.

Several synapse-related terms were also enriched in the prairie vole green and meadow vole darkorchid WGCNA modules which were strongly correlated with the AMY (Fig. 8). Additionally, the terms *cognition* and *behavior* were enriched in both modules, while *learning* and *associative learning* were enriched only in the prairie vole green module. Individual recognition is an essential aspect of bonding behaviors (17), one presumably mediated by these suites of genes. Taken together, these results point to the active role of the AMY in navigating social interactions, with a particular importance in prairie voles for developing social memory for a new partner during bond formation.

The gene categories we found to be enriched in the HT are also consistent with known functions of this region, a diverse structure important for sensing and producing hormonal signals (57). In the contrast between HT and VP/NAc, we found enrichment of the terms *peptide receptor* and *neuropeptide receptor* (Fig. 4f). Similarly, these terms along with several others related to hormone signaling were enriched in the prairie vole turquoise module which was strongly correlated to HT (Fig. 9a). Somewhat surprisingly, we did not find these terms to be enriched in the meadow vole seagreen module which was also strongly correlated with HT (Fig. 9b), though we would expect genes related to hormone and peptide signaling to be just as important in that species as they are in prairie voles. We interpret this as reflecting the well-known role of the hypothalamus in neuroendocrine coordination (57). Another set of GO terms that were enriched in HT compared to both AMY and VP/NAc as well as in both the prairie vole turquoise and meadow vole seagreen modules were several terms related to cilia (Fig. 4d, f; Fig. 9). These genes are likely related to the ependymal cells that lie along the surface of the ventral part of the third ventricle, which is flanked on either side by the HT (58). These cilia play an important role in regulating the flow of cerebrospinal fluid through this ventricle. Other groups of terms that were enriched in the HT and related modules included several related to the endoplasmic reticulum and the extracellular region. The endoplasmic reticulum is involved in protein synthesis, including of neuropeptides (59), so enrichment of this category may be related to neuropeptide synthesis and release.

Many of the terms enriched in the VP/NAc compared to either the AMY or HT (Fig. 4 e, f)—or in the meadow vole gold module (Fig. 10), which was strongly correlated to VP/NAc—were related to the nucleus, chromosomes, and transcription/translation. In addition, there were several enriched terms related to synaptic function including *postsynapse* and terms related to dendritic spines in both the comparison with the HT and in the meadow vole gold module and the term *regulation of postsynaptic neurotransmitter receptor internalization* in the gold module. Together, these enriched terms paint a picture of a transcriptionally actively region with a potentially important role for the formation new synapses or modulation of existing ones.

### Post-mating gene expression in support of bond formation

Pair-bond formation is a major life history transition that is dependent on mechanisms of social reward and memory (17). The necessity of these systems in bonding is the reason we chose to focus on the AMY, HT, and VP/NAc as regions of interest. Prior studies have shown that the AMY is necessary for social recognition and encodes valence (22). The HT is a diverse structure that has several functions related to social behavior (60). Finally, the VP/NAc form a circuit important for action selection. Both dopaminergic and OT inputs to NAc are necessary for pair-bond formation in prairie voles (31, 32). Activation of the D2 dopamine receptor in NAc reduces activity of inhibitory neurons in that region, which is proposed to then disinhibit VP neurons which guide behavioral responses to rewarding stimuli, such as mating (17, 61).

To better understand the molecular mechanisms at work in these regions in response to mating, and during pair-bond formation in prairie voles, we first focused on the genes in each region that were most significantly differentially expressed (lowest FDR) in our model testing for the effect of species:time interaction. In the AMY, this was *Npas4*, which peaked in expression for both species at the 0.5 hr post-mating time point but was expressed at higher levels in prairie voles pre-mating and at later post-mating time points (Fig. 5d). *Npas4* is a transcription factor that plays an important role in the development of inhibitory synapses in response to excitatory inputs by regulating expression of activity-dependent genes (37). In mice, social encounters drive increased *Npas4* expression in the hippocampus and knock out of *Npas4* results in alterations of social behavior, including reduced time investigating novel conspecifics (62). Knockout animals in that study also showed deficits in learning and memory tasks.

Mating is a highly-socially salient behavior that increases neural activity in the AMY (63) so it is unsurprising that *Npas4* peaks in expression shortly after mating onset. However, the fact that it differs across species both before mating, and at later time points following mating onset suggest that it may play a particularly important role for AMY function in prairie voles compared to meadow voles. It may be that AMY activity in prairie voles is simply higher in the absence of mating or other social cues, driving *Npas4* expression more strongly. Alternatively, it may be expressed at constitutively higher levels in prairie voles in the absence of strong activation so that neurons in the region are prepared to develop inhibitory synapses in response to strong excitatory inputs. Either way, this result points to the importance of maintaining excitation/inhibition balance in the AMY for prairie voles.

In the HT, the most strongly differentially expressed gene across species over post-mating time points was *Aqp3* (Fig. 5e), a gene encoding a water/glycerol transporting channel (38) that was lowly expressed in both species until the 12 hr time point, where it increased in expression in meadow voles but not prairie voles. In rats, *Aqp3* is expressed in astrocytes and neurons of several regions, and weakly expressed in ependymal cells, but no expression was found in either the supraoptic or suprachiasmatic nuclei of the hypothalamus (64). Expression of *Aqp3* increased in response to stroke in rats (65) and is elevated in pyramidal cells of cortical tissue from human patients with edema (66); however, the reason why this gene increases in expression in the HT of male meadow voles 12 hrs after beginning to mate with a novel female is not immediately clear.

The gene that was most strongly differentially expressed in the VP/NAc between species across post-mating time points was *Spata16* (Fig. 5f). This gene showed an interesting opposing pattern of expression across species, starting at high levels and reducing quickly following mating in prairie voles and beginning at low levels of expression and peaking shortly after mating in meadow voles. *Spata16* is a gene important for spermatogenesis (43), and to our knowledge it has not so far been studied for its function or localization in the brain.

In addition to the genes most strongly differentially expressed in our model testing the effect of the species:time interaction, we found four genes that significantly differed across species in multiple brain regions (Fig. 6). Among these were *Nfkbia*, which encodes an inhibitor of the transcription factor nuclear factor kappa B (67); *Pxn*, which acts as an adaptor protein that interacts with the cytoskeleton and signaling molecules that regulate cell adhesion to the extracellular matrix (68); the circadian clock gene *Per1* (41); and *Cables1*, which encodes a protein that promotes neurite outgrowth in neurons and plays a role in regulating cell death (69).

Interestingly, though the absolute level of expression varied across regions within a species, the pattern of expression of these genes was largely similar across our focal regions. This suggests that whatever function they serve in response to mating is consistent across the brain regions. Further, these genes tended to increase after mating in meadow voles, then return to baseline, while maintaining relatively more consistent expression across all time points in prairie voles. This suggests that they are responsive to mating in the promiscuous species but are not necessarily important for bonding. Alternatively, it is possible that the changes in expression over time we observe in some or all of these genes are related to time itself, rather than time relative to mating as nearly half of protein-coding mammalian genes show circadian variation in their expression (70). If this is the case, their variation may reflect differences in circadian cycles of gene expression across species.

In addition to these genes that were differentially expressed across species, we also examined modules that were significantly correlated with mating status to identify enriched gene categories associated with these modules. The cyan and pink modules were each positively correlated with having mated, and enriched terms in these modules included many related to translation, ribosomes, and mitochondria. Terms related to catabolic and metabolic processes were also enriched in the pink module. The terms enriched in these modules are likely associated with transcriptional and energetic demands associated with the response to a novel social encounter and initial mating activity. For example, an earlier study has found that 30 min of mating elicits activity-dependent expression of the immediate early genes Fos and Egr-1 in dopaminergic neurons of the AMY in male prairie voles (71). Consistent with this interpretation, the post-mating time point these modules were most strongly related to was 0.5 hr (Fig 7a). Cyan was significantly positively correlated with this point (r=0.33, p=0.024) while pink trended toward a significant positive correlation (r=0.25, p=0.088).

Similar to the modules positively associated with mating, the modules negatively correlated with mating (and so positively correlated with virgin status) were most negatively associated with the 0.5 hr post-mating onset time point (Fig. 7a). The blue module was significantly negatively correlated with this time (r=-0.32, p=0.031) and salmon module trended toward a negative correlation (r=-0.26, p=0.073). Many of the terms enriched in the blue module were related to chromosomes and chromatin modifications, suggesting that activity of these genes is reduced in early mating. In the salmon module, most enriched terms were associated with neural development and axon guidance. As the salmon module was positively correlated with HT, this result suggests that during early mating there is a reduction in the formation or growth of axons from neurons in this region. Together, these results suggest that during very early mating in prairie voles, expression of genes involved in synaptic plasticity and activity of epigenetic modifications are actually reduced compared to the virgin state, perhaps due to the energetic and protein synthesis demands occurring at this time. Though it remains possible that the products of these genes are still present and active during early mating.

### Limitations and future studies

One limitation of the current study is the fact that annotations for the genes encoding OT, OT receptor, and the V1a AVP receptor are absent from the currently available annotated prairie vole genome. While their absence is unfortunate, as these genes would have served as useful positive controls, given the fact that they have been a major focus of studies of mating and pair bonding in voles and other taxa (3,7,17,55,72– 78), it is unlikely that the current study would reveal any novel function for these genes. So, although their inclusion in the annotated genome would have been useful, their absence does not impede the key goals of this study.

While this study identified several gene categories that differ between virgin prairie and meadow voles and across regions in both species, as well as intriguing candidate genes that vary over time after mating across species, we are not able to tie these genes and GO categories to specific behaviors. Future studies that include more detailed behavioral observation will be useful for correlating expression of genes of interest to specific mating-related or other prosocial behaviors that voles engage in during bond formation. Additional manipulative studies will be useful in determining which gene expression differences between prairie and meadow voles.

## Conclusions

Our results emphasize the importance of pre-mating differences in gene expression that confer the ability to pair bond in prairie voles but not in non-bonding species such as meadow voles. In addition, they support the hypothesis that pair-bond formation relies on transcriptional regulation and alterations of neuronal structure in regions including the amygdala, hypothalamus, ventral pallidum, and nucleus accumbens that links neural encoding of partner cues with the reward system, resulting in reinforcement of the partner that leads to selective affiliation. Finally, we identify several intriguing candidate genes that may play important roles in bond formation following co-habitation and mating. Together, our results should help broaden the scope of research on the neural and molecular mechanisms of pair-bond formation by providing novel candidates to investigate in future manipulative studies.

## Methods

### Animals

All animals used in this study were taken from laboratory breeding colonies at Emory University, derived from field-captured voles collected in Illinois. Prairie voles and meadow voles were housed separately in same-sex groups of two to three voles per cage from postnatal day 21. Housing consisted of a ventilated 36cm x 18cm x 19cm plexiglass cage filled with Bed-o’-Cobs laboratory animal bedding (The Andersons Inc., Maumee, Ohio) under a 14/10 hour light/dark cycle (lights on 7:00AM-9:00PM) at 22°C with access to food (rabbit diet; LabDiet, St. Louis, Missouri) and water *ad libitum*. All procedures were approved by the Emory University Animal Care and Use Committee.

### Tissue Collection

All animals were 60-90 days old and sexually naïve at the time of the experiment and tissues were collected from males only. Tissues for the virgin group (time point 0) were collected from sexually naïve males without exposure to females (n=4 per species). For tissues from mated animals, single male voles were paired with a single unrelated female of the same species primed with estradiol benzoate (Sigma Aldrich, St. Louis, MO, USA, BP958) for two days prior to pairing to ensure sexual receptivity. Pairs were observed and the time of first intromission was recorded. Males were collected at 30 minutes (n=4 per species), 2 hours (n=4 per species), and 12 hours (n=4 per species) following the first intromission. Following collection, animals were euthanized using isoflurane, and whole brains were harvested and dissected on a block of dry ice. The AMY, HT, and VP/NAc were collected from each brain and stored in RNAlater (Applied Biosystems AM7020).

### RNA extraction and sequencing

Total RNA was extracted from each region from all individuals using TRIzol (Sigma-Aldrich) following manufacturer’s protocol. An additional DNAse treatment (Purelink) was included to eliminate any contaminating DNA. RNA quantity and quality were assessed using a BioAnalyzer (Agilent) and quantified by spectrophotometry. cDNA was prepared using Superscript reverse transcriptase II (Invitrogen) and purified with QIAquick PCR purification kit. After checking cDNA quantity and quality in the BioAnalyzer, libraries were prepared using Illumina’s TruSeq Sample Prep Kit with starting amount of 1.25µg cDNA. Libraries were normalized using the Evrogen Trimmer kit (Evrogen). Libraries were sequenced as 50 bp single-end reads at the Emory University genomics core facility using Illumina GA_IIX_, HiScan, and HiSeq 1000 and 2000 machines. Read files were trimmed and cleaned using fastq tools.

### Differential gene expression

Reads were aligned to an annotated prairie vole genome (MicOch1.0, INSDC Assembly GCA_000317375.1, Full genebuild annotation by Ensembl released February 2017) using STAR aligner (v2.7.2b) (79). The R package arrayQualityMetrics (80) was used to assess sample quality and identify outliers. One sample was removed for having low read counts and two removed as outliers resulting in 47 prairie vole (n=16 AMY samples, n=15 HT samples, n=16 VP/NAc samples) and 46 meadow vole (n=16 AMY samples, n=15 HT samples, n=15 VP/NAc samples) samples. Genes with less than one count per sample on average were removed prior to analyses.

The dataset was characterized and differential gene expression was analyzed with the R package DESeq2 (34) following the workflow provided by the package authors (81). For analyses of the full dataset, a DESeq dataset was created with the model ∼species + region + time. Poisson distance was calculated using the PoiClaClu package (33). Variance stabilizing transformation and principal component analysis (PCA) were conducted using DESeq2 functions. A likelihood ratio test with the reduced model ∼species + time was used to test for genes that were differentially expressed across regions. To identify genes that were differentially expressed across species before mating, the full model ∼species + region was applied to count data from both species from the virgin time point only, and the likelihood ratio test was used with the reduced model ∼region. To test for species differences in gene expression within a region following mating, the data were subset by region and a likelihood ratio test was used with the full model ∼species + time + species:time and reduced model ∼species + time. This model tests whether species differ in gene expression at any time point after time 0 (virgin animals) (81). Additional annotation information for genes of interest (Tables 1-3) was obtained from the UniProtKB (uniprot.org) manually annotated, reviewed (Swiss-prot) database. Manually annotated gene IDs and descriptions were derived from mouse (*Mus musculus*) annotations.

### Gene correlation networks

We used the R package WGCNA (44) to identify networks of genes with correlated expression. Count data were divided by species, and separate networks were created for prairie voles and meadow voles. Lowly expressed genes (mean of less than 10 counts per sample) were removed and count data were transformed using DEseq2 variance-stabilizing transformation (34). We identified signed modules (i.e. modules consist of genes with strong positive correlation to one another) of correlated genes within networks for each species and found correlations between module eigengenes and sample traits including brain region, mating status (coded 1 for any post-mating time point, 0 for virgin), time of collection (0 [virgin],0.5, 2, or 12 hrs), and individual post-mating collection time points. Following this, we compared species networks by calculating module membership score (kME) (44) for each gene in each module for each species, then merged the kME datasets. We used R package pheatmap (82) to create heatmaps based on module distances and Pearson correlations and used hierarchical clustering by the function hclust with method “average” to visualize relationships of modules across species.

### Gene ontology term enrichment

To test for enrichment of GO terms among genes associated with our differential expression comparisons or WGCNA modules of interest, we used the Mann-Whitney U (MWU) test (35) implemented by the GO_MWU R script (36) (Available at https://github.com/z0on/GO_MWU). Rather than test for enrichment among genes determined to be significant by an FDR cutoff, the MWU test determines if GO categories are enriched using continuous measures. For tests of enrichment among differential expression comparisons, we used signed -log_10_ of the p-value as the measure of comparison (36). The -log_10_ of the p-value from the model was calculated for each gene, then “signed” by multiplying by -1 if the log_2_ fold change (LFC) for the contrast of interest was negative (36). For tests of enrichment in WGCNA modules of interest, we used gene kME values for genes included in the module, and all genes not included in the module were assigned a value of zero and adjusted p-value was determined by 100 permutations where significance measures are randomly shuffled among genes (36).

### Data analysis and visualization

All analyses were conducted using R (v4.0.0). Details of packages and scripts used for differential expression, gene correlation network, and GO term enrichment analyses are described in the relevant sections above. Plots were made using DESeq2 (34) and WGCNA (44) as well as the packages ggplot2 (83), pheatmap (82), and RColorBrewer (84).

## List of abbreviations

AMY: Amygdala
AVP: Arginine-vasopressin
FDR: False discovery rate
HT: Hypothalamus
LFC: Log (base 2) fold change
ME: Module eigengene
MWU: Mann-Whitney U test
OT: Oxytocin
VP/NAc: Ventral pallidum/nucleus accumbens

## Declarations

### Ethics approval and consent to participate

All experiments were approved by the Institutional Animal Care and Use Committee of Emory University.

### Availability of data and materials

The RNA-sequencing reads supporting the results of this article are available in the NCBI Sequence Read Archive under BioProject ID PRJNA682808.

### Competing interests

The authors declare that they have no competing interests

### Funding

This project was supported by National Institutes of Health grants RC1GM090950 to JWT and LJY, P50MH100023 to LJY and ORIP/OD P51OD011132 to Yerkes National Primate Research Center. LAM’s contribution was supported by 1F32MH079661. Additional support was provided by the Intramural Research Program of the National Human Genome Research Institute, National Institutes of Health to JWT.

### Authors’ contributions

L.A.M. and L.J.Y. conceived the study. L.A.M. collected tissue, J.K.D. prepared samples for sequencing, J.W.T. oversaw sequencing. A.B., J.A.T., M.M. and S.M.P. analyzed the data. A.B., J.A.T., and S.M.P. wrote the manuscript. All authors approved of the final manuscript.

## Acknowledgements

The authors thank Aubrey M. Kelly and Annaliese Beery for providing vole images.

## Notes

### Competing Interest Statement

The authors have declared no competing interest.

